# Cell type specificity of mosaic chromosome 1q gain resolved by snRNA-seq in a case of epilepsy with hyaline protoplasmic astrocytopathy

**DOI:** 10.1101/2023.10.16.562560

**Authors:** Kun Leng, Cathryn R. Cadwell, W. Patrick Devine, Tarik Tihan, Zhongxia Qi, Nilika Singhal, Orit Glenn, Sherry Kamiya, Arun Wiita, Amy Berger, Joseph T. Shieh, Erron W. Titus, Mercedes F. Paredes, Vaibhav Upadhyay

## Abstract

**Introduction:** Mosaic gain of chromosome 1q (chr1q) has been associated with malformation of cortical development (MCD) and epilepsy. Hyaline protoplasmic astrocytopathy (HPA) is a rare neuropathological finding seen in cases of epilepsy with MCD. The cell-type specificity of mosaic chr1q gain in the brain and the molecular signatures of HPA are unknown.

**Methods:** We present a child with pharmacoresistant epilepsy who underwent epileptic focus resections at age 3 and 5 years and was found to have mosaic chr1q gain and HPA. We performed single-nuclei RNA-sequencing (snRNA-seq) of brain tissue from the second resection.

**Results:** snRNA-seq showed increased expression of chr1q genes specifically in subsets of neurons and astrocytes. Differentially expressed genes associated with inferred chr1q gain included *AKT3* and genes associated with cell adhesion or migration. A subpopulation of astrocytes demonstrated marked enrichment for synapse-associated transcripts, possibly linked to the astrocytic inclusions observed in HPA.

**Discussion:** snRNA-seq may be used to infer the cell type-specificity of mosaic chromosomal copy number changes and identify associated gene expression alterations, which in the case of chr1q gain may involve aberrations in cell migration. Future studies using spatial profiling could yield further insights on the molecular signatures of HPA.

## INTRODUCTION

Mosaic chromosome 1 q (chr1q) gain has been identified in several case reports and series as a genetic driver of pharmacoresistant epilepsy in the setting of malformation of cortical development (MCD)^1,2,3,4,5^, an umbrella term which encompasses pathologies such as mild MCD (mMCD) with excessive heterotopic neurons^6^, focal cortical dysplasia (FCD), gray matter heterotopia, polymicrogyria, and hemimegalencephaly.

Hyaline protoplasmic astrocytopathy (HPA) is a rare neuropathological finding often associated with epilepsy and MCD^7^ (see eAppendix 1) characterized by protoplasmic astrocytes laden with electron-dense, non-membrane-bound hyaline inclusions reported to have filamin A, GLT-1 (SLC1A2), α-β-crystallin, and cytoglobin immunoreactivity^8,9^.

Here we present a child with pharmacoresistant epilepsy who was found to have mosaic chr1q gain as well as HPA from two sequential epileptic focus resection surgeries. We performed single-nuclei RNA-sequencing (snRNA-seq) of brain tissue from the second resection to gain insights into the cell type-specificity of his mosaic chr1q gain and the molecular signatures associated with HPA.

## METHODS

### Research ethics and informed consent

Tissue was acquired and deidentified by the Neurosurgery Tissue Bank at the University of California, San Francisco with patient consent in strict observance of the legal and institutional ethical regulations under the UCSF Committee on Human Research (IRB # 10-01318). The tissue was snap frozen in liquid nitrogen in less than 30 minutes after surgical resection.

### Single-nuclei RNA-sequencing (snRNA-seq)

snRNA-seq was performed on fresh frozen brain tissue from the patient’s second epileptic focus resection. 128 mg of frozen brain tissue was cut at –20 °C and sent on dry ice to SingulOmics Corporation (New York) for nuclei extraction and snRNA-seq using the 10x Genomics v3 platform, targeting recovery of 10,000 nuclei and 25,000 reads per cell. ∼200 million PE150 reads were sequenced on an Illumina NovaSeq 6000 sequencer.

### Data analysis

Processing and mapping of raw reads from snRNA-seq and cell calling were performed with Cell Ranger (v7.1.0) using GRCh38-2020-A with inclusion of intronic regions as the reference genome. Analysis of snRNA-seq data was performed in R (v4.3) using Seurat (v5). See eMethods in Supplemental Information.

### Data availability

Raw reads and the filtered feature-barcode matrix from snRNA-seq are available on GEO (GSE241521).

## RESULTS

### Clinical course

Our patient presented with seizures at 6 months of age. For details of his initial clinical presentation and workup, see eAppendix 2. 3T MRI at age 40 months demonstrated very subtle increased T2-FLAIR signal throughout the white matter of the anterior right frontal lobe with minimal blurring of the gray-white border, raising suspicion for FCD (Fig. 1a), which was not visible on 3T MRIs done at age 6 months and 23 months due to incomplete myelination of the frontal lobes. PET scan at age 40 months showed right frontal lobe hypoperfusion (Fig. 1a).

**Fig. 1.**
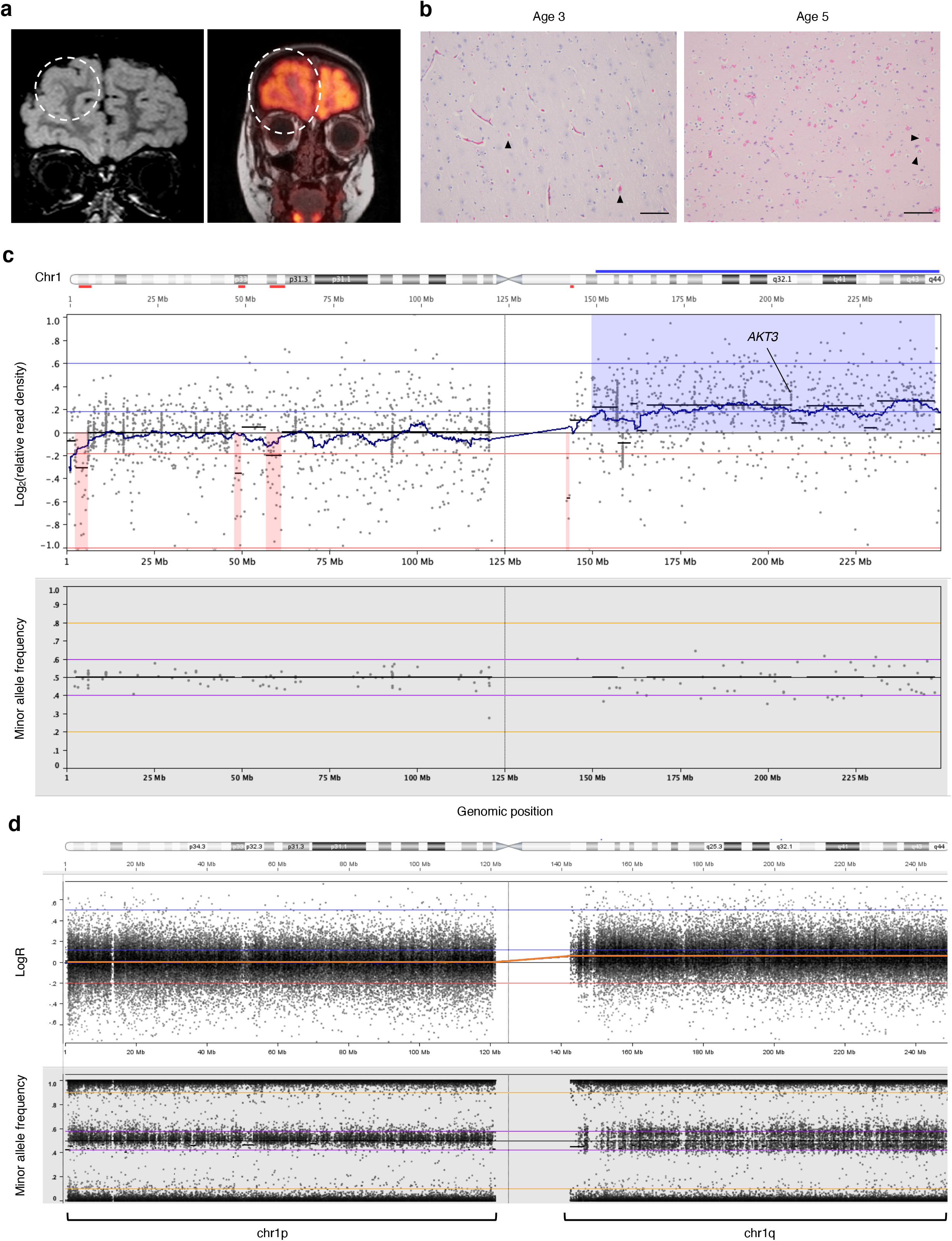
Right frontal lobe abnormalities on neuroimaging, hyaline protoplasmic astrocytopathy on neuropathology, and evidence of mosaic chr1q gain on genomic analyses. **a**, 3T T2-FLAIR MR image (left) at age 40 months demonstrating subtle increased hyperintensity in the right frontal white matter with minimal blurring of the gray-white border (dotted circle) suggestive of focal cortical dysplasia; PET image (right) at age 3 years demonstrating hypoperfusion in the right frontal lobe (dotted circle). **b**, H&E sections of surgically resected cortical tissue from the surgeries at age 3 and 5 years demonstrating worsened hyaline protoplasmic astrocytopathy (representative inclusion-laden astrocytes labeled by black arrowheads) over time. Scale bar, 100 µm. **c**, Visualization of relative read density (top) and minor allele frequency of common single nucleotide polymorphisms (bottom) from the UCSF 500 Cancer Gene Panel performed on surgically resected brain tissue from the surgery at age 3 years. Regions highlighted in blue in the top subpanel reflect regions of chr1q with definitive copy number gain; the jagged appearing black line in the top subpanel corresponds to the running average of the relative read density, and the horizontal black lines in the top subpanel correspond to the average relative read density within corresponding genomic regions. **d**, Visualization of LogR values (see eMethods) and minor allele frequency derived from the Illumina CytoSNP 850 K array applied to genomic DNA extracted from neuronal nuclei isolated by fluorescence-activated nuclei sorting from surgically resected brain tissue at age 3 years. The orange line reflects the running average of the LogR value along chr1. The remaining colored horizontal lines serve as visual guides.

An intraoperative electrocorticography (ECOG)-guided right frontal lobe resection was performed at age 40 months to remove the epileptogenic focus (eFigure 2a). On neuropathology, HPA was seen (Fig. 1b; eFigure 2b). In addition, there appeared to be an abnormal number of heterotopic neurons (> 30/mm^2^) in the subcortical white matter (eFigure 2c) without definite cortical dysplasia (eFigure 2b), consistent with the new ILAE classification of mMCD with excessive heterotopic neurons^6^. Genomic analysis of the resected brain tissue using the UCSF500 Cancer Gene Panel with a buccal swab sample for comparison revealed an approximate 1.2-fold gain in copy number of chr1q in the resected brain tissue (Fig. 1c) but not in the buccal sample, indicative of mosaic chr1q gain in the brain. Due to continued seizures and failure of medical management, a motor-sparing right frontal lobectomy was performed at age 5 years. The resected tissue from the second surgery demonstrated an approximate 1.05-1.1-fold gain in chr1q copy number, which was lower compared to prior, but with more extensive HPA (Fig. 1b). Electron microscopy performed on tissue from the second surgery showed numerous large inclusions composed of electron-dense, non-membrane-bound granular material (eFigure 2d). The inclusions appeared to be predominantly in cortical astrocytes and were frequently juxtanuclear in location, consistent with prior ultrastructural studies of HPA^8,9^.

### Transcriptomic profiling of surgically resected brain tissue

We performed snRNA-seq of fresh frozen brain tissue from our patient’s second surgery, recovering nuclear transcriptomes corresponding to 10,764 cells at ∼23,000 mean reads per cell and 1,880 median genes detected per cell. After initial clustering and quality control (see eMethods), 10,179 cells were retained for further analysis. Uniform manifold approximation projection (UMAP) of above cells showed well-separated clusters (Fig. 2a) of excitatory and inhibitory neurons, astrocytes, oligodendrocytes, oligodendrocyte precursor cells, microglia, endothelial cells, and vascular cells, which were assigned their respective cell types based on alignment to a gold-standard reference dataset (see eMethods). All clusters demonstrated expression of appropriate cell type-specific genes (eFigure 3a) and acceptable quality metrics (eFigure 3b).

**Fig. 2.**
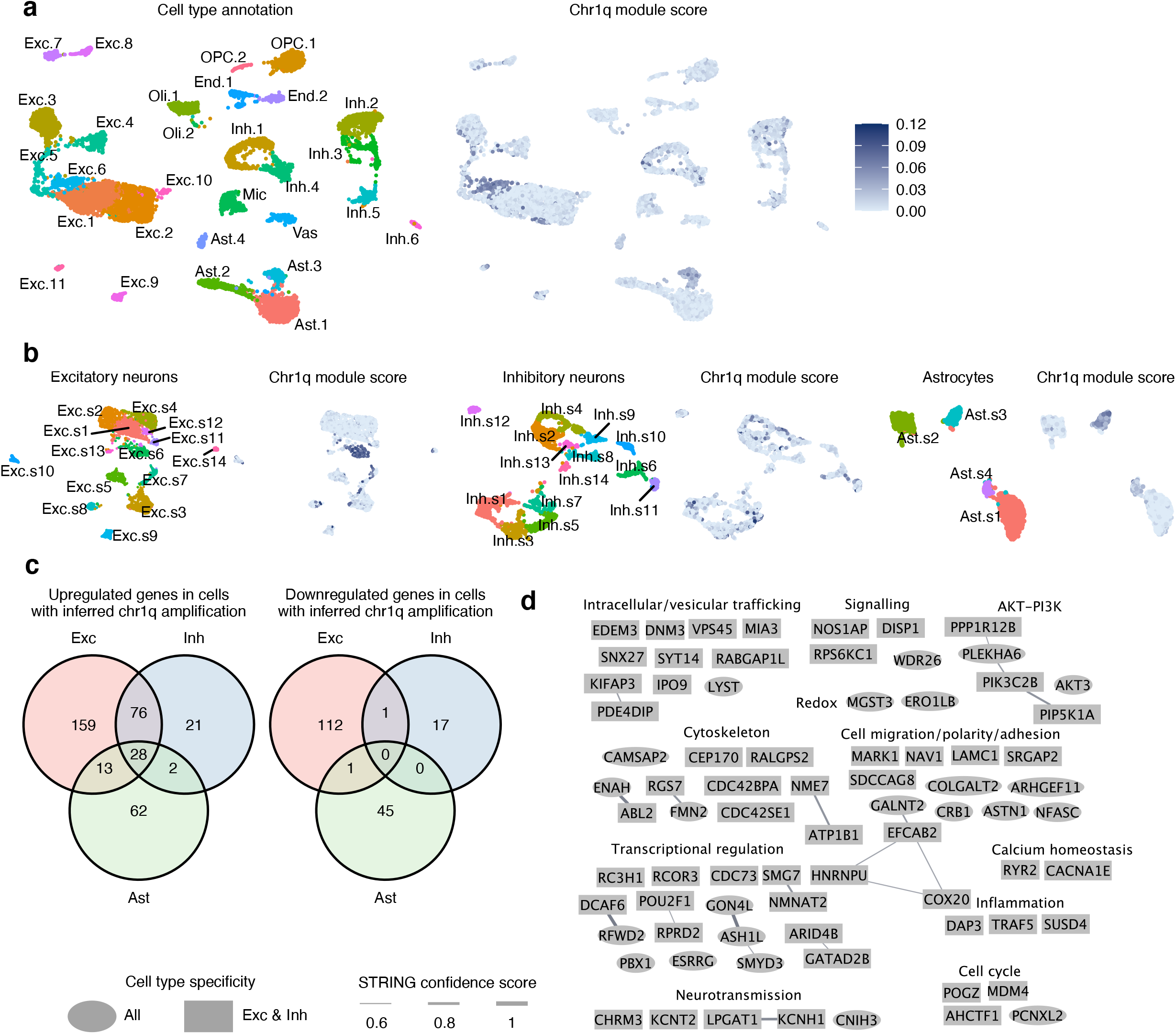
Mosaic chr1q gain in neurons and astrocytes and associated alterations in gene expression identified by snRNA-seq. **a**, Uniform manifold approximation projection (UMAP) of cells recovered from snRNA-seq, colored by cell cluster assignment (left) or chr1q module score (right). **c**, UMAP of excitatory neuron, inhibitory neuron, or astrocyte subclusters colored by subcluster assignment or chr1q module score. **c**, Venn diagrams of upregulated (left) or downregulated (right) differentially expressed genes (DEGs) associated with chr1q gain in excitatory neurons, inhibitory neurons, or astrocytes. **d**, Network visualization of chr1q gain associated upregulated DEGs shared between excitatory and inhibitory neurons grouped by function. Lines connecting genes reflect evidence of regulatory relationships (see eMethods). Exc – excitatory neurons, Inh – inhibitory neurons, Ast – astrocytes, Oli – oligodendrocytes, OPC – oligodendrocyte precursor cells, Mic – microglia, End – endothelial cells, Vas – vascular cells.

To infer the cell type-specificity of mosaic chr1q gain, the expression of genes on chr1q was compared to that of all other genes to generate a chr1q module score (Fig. 2a-b, eFigures 4-5). In parallel, we also employed inferCNV to conduct an unbiased assessment of chromosomal copy number (eFigure 6, see eMethods). Both methods revealed chr1q gain to be restricted to neurons and astrocytes (eFigures 4-6). See eAppendix 3 for a detailed analysis. Next, we experimentally confirmed mosaic chr1q copy number gain in neurons using fluorescence-activated nuclei sorting followed by RT-PCR (see eMethods and eFigure 7) and high-density SNP array analysis (Fig. 1d, eFigure 1b) on genomic DNA extracted from sorted nuclei.

Differentially expressed genes (DEGs) in cells with inferred chr1q gain compared to those without (see eMethods) showed cell type-specificity, with 28 upregulated genes shared among all three cell types (Fig. 2c, eFigure 8, eTable 1). Among the shared DEGs were *LYST*, a regulator of lysosomal and endosomal trafficking, and *AKT3*, an upstream activator of the mTOR pathway; both genes are located on chr1q. Activating somatic mutations in *AKT3* have been associated with epilepsy and MCD and may drive pathology through overactivation of the mTOR pathway^10,11^. However, we did not recover a significant enrichment for the mTOR pathway in chr1q gain associated DEGs, which could be attributed to the fact that mTOR acts mainly through its kinase activity and does not directly affect transcription. Chr1q gain associated DEGs demonstrated enrichment for genes involved in cellular adhesion and intercellular junctions (eFigures 9-12, eTable 2), processes which are important for cell migration. Given that cortical development requires the appropriate migration of neuronal progenitors and young migratory neurons, we suspected that chr1q gain associated DEGs in neurons may be involved in cellular migration. Indeed, examination of NCBI and UniProtKB gene summaries of upregulated DEGs shared between excitatory and inhibitory neurons revealed annotations for processes such as cytoskeletal regulation (e.g. *FMN2*), vesicular trafficking, cellular adhesion, and cellular migration (e.g. *ASTN1*); furthermore, there was evidence of regulatory interactions among many of these genes (Fig. 2d).

We next turned our attention to evaluate whether chr1q gain exhibited enrichment in cell type subpopulations by subcluster analysis excitatory neurons, inhibitory neurons, and astrocytes. We found subclusters with marked enrichment of chr1q-gained cells in excitatory neurons (Exc.s6) and astrocytes (Ast.s3, Ast.s4), but not inhibitory neurons, where chr1q-gained cells were distributed more evenly across subclusters (Fig. 2b). In excitatory neurons, Exc.s6 compared to all other excitatory neurons demonstrated differential expression of genes involved in neuronal projections (eFigure13, eTables 3-4). In astrocytes, Ast.s3 compared to all other astrocytes demonstrated downregulation of genes involved in glutamate uptake, such as *SLC1A2*, and potassium transport, such as *KCNIP4* (eFigure 14, eTable 3-4). Notably, Ast.s4 compared to all other astrocytes demonstrated a striking enrichment of synapse-associated transcripts such as *SHISA9*, *GRIA1*, *GRIN2A*, *GRIN2B*, *NGLN1*, and *NRXN3* ^12^ in conjunction with upregulation of *HSP90AA1* (Fig. 3a-c, eFigure 15, eTables 3-4), which encodes the Hsp90 chaperone. We confirmed the presence of this astrocyte subpopulation in a recently published snRNA-seq dataset of additional patients with mosaic chr1q gain and HPA (Miller *et al.*)^13^ (eFigure 16).

**Fig. 3.**
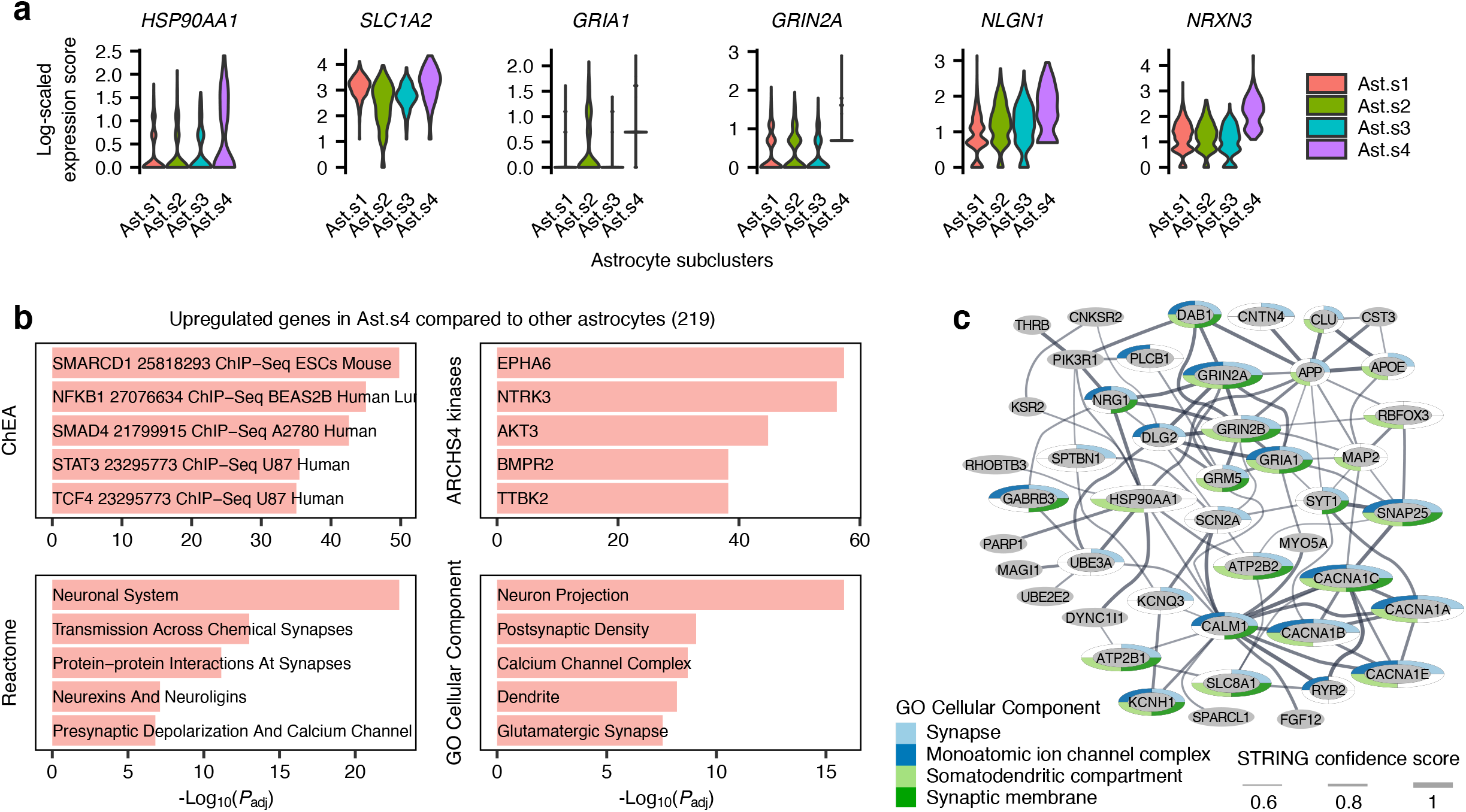
A subpopulation of astrocytes enriched in synapse-associated transcripts identified on subclustering analysis. **a**, Log-scaled expression of selected synapse-associated transcripts in astrocyte subclusters. **b**, Enriched terms from the ChEA, ARCHS4, Reactome, and GO Cellular Component gene set libraries (see eMethods) for genes upregulated in Ast.s4 compared to other astrocytes. **c**, Network visualization of upregulated genes in Ast.s4 that have known regulatory relationships with Hsp90 (*HSP90AA1*) or with those that have a known regulatory relationship with Hsp90.

## DISCUSSION

The enrichment of chr1q gain associated DEGs for genes involved in cell adhesion and migration, which was also seen in Miller *et al.*^13^, suggests that chr1q gain disrupts the developmental migration of neuronal and glial progenitor cells. See eAppendix 4 for further discussion.

In astrocytes, chr1q gain may disrupt homeostatic astrocyte functions such as glutamate uptake and potassium buffering (as seen in Ast.s3), which may contribute to epilepsy. Furthermore, we discovered a subpopulation of astrocytes with overrepresentation of chr1q-gained cells and marked enrichment of synapse-associated transcripts (Ast.s4), which was also present in the data from Miller *et al.*^13^ (eFigure 16). We hypothesize that the enrichment of synapse-associated transcripts in these astrocytes stems from phagocytosis of synaptic debris^14^. This hypothesis is supported by the fact these astrocytes have increased expression of *ADGRB1* (eFigure 16f), which encodes BAI1, an adhesion GPCR that recognizes phosphatidylserine^15^. We speculate that with cytoskeletal, vesicular trafficking, and lysosomal abnormalities secondary to chr1q gain, these astrocytes may have exceeded their degradative or intracellular trafficking capacity, which may contribute to the inclusions observed in protoplasmic astrocytes on neuropathology and electron microscopy. Please see eAppendix 5 for additional discussion.

Lastly, please see eAppendix 6 for additional discussion of the clinical findings and eAppendix 7 for additional discussion of the methodological merits and limitations of our snRNA-seq results.

In summary, we identified cell type-specific changes in gene expression associated with chr1q gain and possibly HPA in our patient concordant with findings in similar patients, which pointed towards cellular pathways potentially affected by these pathologies. Ultimately, spatial transcriptomics and proteomics will further elucidate the molecular signatures and dysregulated cellular functions associated with chr1q gain and HPA.

## AUTHOR CONTRIBUTIONS

KL conducted the experiments, performed the data analysis, and wrote the paper, with close input from VU, MFP, and EWT. EWT conceived the project with MFP and VU, led the initial research efforts with assistance from AB and ZQ, and secured funding for the snRNA-seq experiment from the UCSF Department of Laboratory Medicine. NS, JS, TT, OG, and WPD were involved in the clinical care and workup of our patient. CRC contributed to the interpretation of HPA neuropathology. SK assisted with electron microscopy. AW oversaw preparation of the Illumina SNP array and analyzed the data from the array.

## Supporting information

eTable1

eTable2

eTable3

eTable4

## ACKNOWLEDGMENTS

We thank Dr. Alyssa Reddy, Dr. Danilo Bernardo, and Dr. Kurtis Auguste for their involvement in the clinical care of our patient. We thank Dr. Biswa Ramani for assistance with neuropathology interpretation. We thank Julia S. Chu for assisting with storage of clinical samples. We thank Melissa Chow and Todd Johnson for processing the Illumina SNP array. We thank Dr. Clifford Lowell, chair of Laboratory Medicine at UCSF, for funding support. KL was supported by NIA F30AG066418. EWT was supported by Damon Runyon Fellowship DRG122-22. MFP was supported by the Roberta and Oscar Gregory Endowment in Stroke and Brain Research, Chan Zuckerberg Biohub, and NINDS 1R21NS123461-01A1. VU was supported by NHLBI K08HL165106. CRC was supported by NINDS K08NS126573 and U01NS132353, the Weill Neurohub, the Shurl and Kay Curci Foundation, and Citizens United for Research in Epilepsy (CURE).

## eAPPENDICES

**eAppendix 1**

HPA has also been associated with Aicardi syndrome^1^ and tuberous sclerosis^2^, both of which tend to present with refractory epilepsy. In addition to patients with refractory epilepsy, it has also been seen in older patients with dementia^34^.

**eAppendix 2**

Our patient’s seizures were characterized by left-sided limb stiffening and gaze deviation, occurring up to one hundred times a day. Scalp EEG revealed right frontal localization, and MRI at 6 months showed non-specific findings without evidence of cortical malformation. The seizures were resistant to standard anti-seizure medications including levetiracetam, oxcarbazepine and clobazam. GeneDx Infantile Epilepsy panel and whole-exome sequencing did not reveal any pathogenic variants. Suppression of seizures was achieved for one year on vigabatrin, with recurrence of seizures on weaning vigabatrin and no response to subsequent medical management. After the second surgery, our patient’s seizure frequency improved to approximately once every 10 days on a two-drug anti-seizure medication regimen (clobazam, lamotrigine) continued from prior to the surgery.

**eAppendix 3**

We found that chr1q module scores were bimodally distributed, with 10% of cells exceeding the cutoff value of 0.025, which is the chr1q module score corresponding to the local minimum between the two modes (eFigure 4a). In contrast, a module score generated with genes from chr1p as a negative control was unimodally distributed (eFigure 4b). Thus, assuming that cells with chr1q module score > 0.025 correspond to cells with chr1q gain and that cells with chr1q gain are trisomic for chr1q, the above estimate of 10% mosaicism would correspond to a bulk fold-change in chr1q copy number of 1.05, consistent with that derived from analysis of the UCSF500 panel data. Plotting the distribution of chr1q module score for each cell type showed bimodality only in excitatory neurons, inhibitory neurons, and astrocytes (eFigure 4c), with a Pearson’s Chi-squared test *P* value of < 2.2 × 10^-16^ (eFigure 4d), suggesting that chr1q gain is restricted to neurons and astrocytes. To support the above analysis, we used an alternative computational approach, inferCNV^5,6,7,8^, to infer genome-wide copy number changes based on observed transcript levels (see eMethods), which revealed copy number gain in chr1q only in neurons and astrocytes (eFigure 6). We believe that the chr1q gain in our patient is likely trisomic given that cells from our patient exhibit smaller chr1q module scores (e.g. eFigure 4a) compared to cells derived from patients described in Miller *et al.*^9^ (eFigure 16c), where four out of five patients were confirmed to have tetrasomic mosaic chr1q gain.

**eAppendix 4**

Disruption of the developmental migration of neuronal and glial progenitor cells is supported clinically in our patient by white matter T2-FLAIR hyperintensity with blurring of the gray-white border in the right frontal horn on MRI and appearance of increased heterotopic neurons in subcortical white matter on neuropathology. Although excitatory neurons and inhibitory neurons are traditionally thought to arise from distinct pools of precursor cells originating from different regions of the developing brain^10^, the presence of chr1q gain in both excitatory and inhibitory neurons (as well as astrocytes) in our patient is in line with the recent finding that a subset of cortical progenitor cells can potentially give rise to both excitatory and inhibitory neurons during human brain development^11^. The hypothesis that chr1q gain occurred in a cortical progenitor cell during neurodevelopment would also be consistent with *in vitro* data showing that chr1q gain confers a proliferative advantage in neural stem cells^12^. Lastly, the existence of a subpopulation markedly enriched in chr1q-gained cells in excitatory neurons (Exc.s6) but not inhibitory neurons suggests that migration defects may be more pronounced in excitatory neurons compared to inhibitory neurons.

**eAppendix 5**

Ast.s4 has a lower average number of genes recovered per cell compared to other astrocyte subclusters (eFigure 5b) and is less enriched for chr1q-gained cells compared to Ast.s3 (Fig. 2b; 61/122 chr1q-gained cells in Ast.s4 vs. 207/226 chr1q-gained cells in Ast.s3). We believe that the lower observed enrichment for chr1q-gained cells in Ast.s4 could be attributed to the lower number of genes recovered per cell in Ast.s4, which in turn may stem from lower quality snRNA-seq library generation due to the proposed presence of juxtanuclear HPA inclusions in these cells. Interestingly, astrocytes from Miller *et al.* which mapped to Ast.s4 on label transfer analysis also had a lower number of genes recovered on average compared to other astrocytes in Miller *et al.* Thus, it is possible that the true representation of chr1q-gained cells in Ast.s4 is higher than observed.

**eAppendix 6**

The lower degree of mosaicism of chr1q gain from the second surgery compared to the first may reflect a correspondence between the degree of chr1q gain mosaicism and anatomical proximity to the epileptic focus. A similar hypothesis has been validated in the setting of FCD and the rate of mosaic somatic mutations in mTOR pathway genes^13^.

**eAppendix 7**

Our analysis demonstrates that snRNA-seq of archived fresh frozen brain tissue could be used to infer the cell type-specificity of mosaic chromosomal copy number variations and their associated alterations in gene expression, which has been underutilized in neurology research compared to oncology^5,6,7,8^. Thus, single-cell transcriptomics provides a powerful complementary approach to traditional methods such as karyotyping (limited to cells proliferating culture) or FISH. For our patient, tissue culture of freshly isolated primary cells from the second surgery was severely limited by yield, and FISH performed on cultured cells failed to identify any cells with chr1q copy number gain. Per expert consultation, we did not attempt FISH on an FFPE sample due to expected technical challenges (such as cutting artifacts, cell overlap, and unclear cell morphology) that limit sensitivity in the setting of low-level mosaic copy number changes.

## eMETHODS

### Intraoperative electrocorticography

Following a right frontotemporoparietal craniotomy under local anesthesia using nitrous oxide, narcotics, and dexmedetomidine, the cortical surface was exposed. The brain appeared grossly intact. A 5x4 array of 20 electrodes numbered 1 through 20 was placed over the right frontal lobe with the following orientation: 20 - anterior inferior, 1 - posterior superior, 5 - anterior superior, 16 - posterior inferior, motor strip - 1, 6, 11. Recording from these electrodes revealed a background consisting of low-amplitude theta frequency activity with superimposed faster elements.

### Electron microscopy

Freshly harvested tissue was minced into pieces < 2 mm in width. Tissue pieces were fixed with glutaraldehyde in cacodylate buffer and then stored in cacodylate buffer. Osmium tetraoxide was used for secondary fixation, followed by dehydration and resin embedding. Thin sections were stained with uranyl acetate and lead citrate. Images were collected by the UCSF Pathology department electron microscopy service.

### UCSF 500 Cancer Gene Panel

The UCSF 500 Cancer Gene Panel is an exon-capture-based panel that targets 529 genes plus additional genome-wide probes to detect gross chromosomal gains or losses. The test is available to patients through the UCSF Clinical Cancer Genomics Lab (CPT code 81455). For visualization of chromosomal-level copy-number variations, the log_2_ value of the ratio of sample coverage over pooled normal coverage is plotted in Fig. 1c. The pooled normal coverage is derived from ∼300 samples that were processed and sequenced in the same pipeline as the sample. No specific threshold value was chosen to identify genomic copy number changes, given that chr1q clearly had higher relative read density compared to other genomic regions. No other genomic region appeared to have copy number changes.

### snRNA-seq data analysis

The filtered feature-barcode matrix of raw UMI counts generated by CellRanger (v7.1.0) after alignment of raw reads was loaded into Seurat (v5)^14^. Data normalization and identification of highly variable genes was performed using Seurat::SCTransform^15^ with default parameters. The resulting Pearson residuals from SCTransform’s negative binomial generalized linear model correcting for variation in library size among cells (the ‘scale.data’ slot in the ‘SCT’ assay) corresponds to the “log-scaled expression score” used for differential expression analysis and gene expression visualization referenced in the manuscript and described below. Dimensionality reduction and initial clustering was performed using Seurat::RunPCA, Seurat::RunUMAP, Seurat::FindNeighbors, and Seurat::FindClusters with default parameters, specifying a range of values for the resolution parameter for Seurat::FindClusters. After visual inspection of cluster assignments on UMAP, a resolution of 0.8 was deemed satisfactory for initial clustering.

For basic quality control, quality metrics including number of genes detected and percent of genes from the mitochondrial genome (“percent.mito”) was visualized together with cluster assignment on UMAP, identifying a cluster of cells with high percent.mito which was removed. Cluster markers were then identified using Seurat::FindAllMarkers with the following parameters: assay = ‘SCT’, slot = ‘scale.data’, test.use = ‘t’, and the ‘features’ parameter set to the top 3000 highly variable genes from SCTransform. Clusters were then assigned to major known brain cell types (excitatory neurons, inhibitory neurons, astrocytes, oligodendrocytes, oligodendrocyte precursor cells, microglia, endothelial cells, vascular cells) by using cell type predictions generated by Azimuth^16^ (an online analysis platform for alignment of query single-cell datasets to annotated reference single-cell datasets: https://azimuth.hubmapconsortium.org/), using the human primary motor cortex as the reference^17^. A cluster with low mapping scores from Azimuth and lack of cell type-specific markers was removed. After all clusters have been assigned to major known brain cell types, separate seurat objects were created with only cells belonging to each cell type for subclustering analysis.

For subclustering of each cell type, data normalization, identification of high variable genes, dimensionality reduction, and clustering was repeated starting from the raw counts, again specifying a range of values for the resolution parameter for Seurat::FindClusters and using visual inspection of cluster separation on UMAP to select the optimal resolution parameter. Initial cluster markers were then identified using Seurat::FindAllMarkers with assay = ‘SCT’, slot = ‘scale.data’, test.use = ‘t’ and filtered for adjusted *P* value < 0.05 (Bonferroni) and log_2_-fold change > 0.58. Cell type identity of clusters was then verified using Azimuth and inspection of cluster markers. In some cases, this process identified small clusters consisting of cells predicted to be another cell type by Azimuth, likely doublets, which were then removed. The remaining clusters were then assigned as bona fide cell type subclusters. Subcluster markers were then identified using Seurat::FindAllMarkers with the same parameters as above, and filtered for adjusted *P* value < 0.05 (Bonferonni). All cells removed during subclustering were also removed from the seurat object corresponding to overall dataset, which was then used for the analyses described below.

For inference of chr1q gain, the chr1q module score was generated using Seurat::AddModuleScore with assay = ‘SCT’ and the features parameter set to genes located on chr1q, which were identified using Ensembl Biomart and restricted to genes with official HGNC gene symbols. The chr1p module score was calculated similarly.

As an alternative approach to infer presence of chr1q gain, we used inferCNV^5,6,7,8^ to compute relative genomic copy number at the single-cell level, following the instructions available on GitHub (https://github.com/broadinstitute/inferCNV/wiki). The library-size corrected UMI counts from SCTransform (the ‘counts’ slot of the ‘SCT’ assay) was used as input into inferCNV.

To identify chr1q gain-associated DEGs robust to cell type, an expression matrix consisting of Pearson residuals from SCTransform (the ‘scale.data’ slot of the ‘SCT’ assay) of genes that were detected in more than 5% of all cells was used as input to limma::lmFit (limma v3.56.2), using a model formula of the form ‘∼ cell type + chr1q status’. Statistical shrinkage was then performed using limma:eBayes, with robust = ‘TRUE’. DEGs were then filtered for adjusted *P* value < 0.05 (Benjamini-Hochberg) and log_2_-fold change > 0.32. To identify chr1q gain-associated DEGs for each cell type, Seurat::FindMarkers was used with the seurat object corresponding to each cell type, with assay = ‘SCT’, slot = ‘scale.data’, test.use = ‘t’, ident.1 set to cells with chr1q module score > 0.025, and ident.2 set to cells with chr1q module score < 0.025. Since Seurat::FindMarkers uses Bonferonni correction by default, Benjamini-Hochberg adjusted *P* values were generated using p.adjust with method = ‘fdr’, and DEGs were filtered for adjusted *P* value < 0.05 and log_2_-fold change > 0.32. Given that when comparing cells with high chr1q module score to those with low chr1q module score, genes on chr1q are more likely to be differentially expressed, we applied a more stringent *P* value adjustment (Holm’s method) for genes on chr1q.

Enrichment analysis of DEGs was performed for upregulated and downregulated DEGs separately using Enrichr (an online tool for gene set enrichment analysis), utilizing the R utility function enrichR::enrichr with the following gene set libraries: “ENCODE_and_ChEA_Consensus_TFs_from_ChIP-X’, ’ARCHS4_TFs_Coexp’, ’Reactome_2022’, ’ARCHS4_Kinases_Coexp’, ’GO_Biological_Process_2023’, and ’GO_Cellular_Component_2023’. The ENCODE and ChEA consensus library consist of gene sets associated with transcriptional regulators through ChIP-seq experiments. ARCHS4^18^ is a continually updated compendium of all published RNA-seq datasets re-processed through a common pipeline enabling co-expression analysis of genes with known transcription factors (TFs) and kinases. The adjusted *P* values from Enrichr are used for the plots in Fig. S4-6, which show only terms with adjusted *P* value < 0.1.

For network visualization of genes shown in Fig. 2h and Fig. 3c, genes of interest were used as the input to the STRING (v11.5)^19^ app’s “protein query” function in CytoScape^20^ (v3.10.0); a minimum cutoff of 0.6 was used for the confidence score, which reflects the degree of certainty regarding the existence of a regulatory relationship between two proteins based on current evidence. Genes without annotated functions in their NCBI gene summary were not included for visualization.

For label transfer analysis of astrocytes from Miller *et al.*^9^, processed gene-level UMI matrices for each sample were downloaded from the Gene Expression Omnibus under accession GSE221849, aggregated into a single matrix, and then subject to the same analysis pipeline as described above. Astrocytes from Miller *et al.* were then mapped to astrocytes in our manuscript (serving as the reference) using Seurat::FindTransferAnchors (reference.assay = ’SCT’, query.assay = ’SCT’, reference.reduction = ’pca’, k.anchor = 20) and Seurat::MapQuery (reference.reduction = ’pca’, reduction.model = ’umap’).

### Fluoresence-activated nuclei sorting (FANS)

Nuclei were extracted from 240 mg of fresh frozen brain tissue from the second surgery at age 3 years and then subject to FANS as described in Mussa *et al.*^21^. Neuronal nuclei were gated as NeuN+. Gating of Non-neuronal nuclei differed from the strategy described in Mussa *et al.* due to lack of strong PAX6 staining in putative astrocyte nuclei, which was noted to occur for a subset of samples in Mussa *et al*. Non-neuronal nuclei (NeuN-) were gated as follows: astrocytes (PAX6lo, OLIG2-), oligodendrocytes (PAX6lo, OLIG2lo), and OPCs (PAX6hi, OLIG2hi).

### Genomic copy-number RT-PCR

Genomic DNA was extracted from nuclei sorted using the gating strategy described above using the NucleoSpin Blood Mini kit from Machery Nagel (cat. no. 740951.50) following the manufacturer’s protocol. Chr1q copy number was determined by RT-PCR using the hPSC Genetic Analysis kit from StemCell Technologies (cat. no. 07550), which includes proprietary double-quenched Taq-man probes against a region of chr1q, following the manufacturer’s protocol. Genomic DNA yield was unexpectedly extremely low for sorted astrocyte and oligodendrocyte nuclei, which precluded successful copy-number determination using RT-PCR for those populations.

### High-density SNP array

Genomic DNA from sorted neuronal nuclei was processed for analysis by Illumina Global Diversity Array with Cytogenetics-8 according to the manufacturer’s protocol. Data were processed and CNV calls were generated with NxClinical v6.0 (BioNano). CNV calls by SNP array were reported using standard clinical thresholds in the University of California San Francisco Clinical Cytogenetics Laboratory, including annotation of any CNVs□>□500 kb. Interpretation of clinical significance was done according to standard American College of Medical Genetics guidelines^22^. LogR values correspond to Log_2_(R_observed_/R_expected_), where R is the normalized spot intensity on the array.

### Statistics

The Pearson’s Chi-square test or Student’s t test was performed with the functions ‘chisq.test’ or ‘t.test’ respectively in R with default parameters.

## eTABLE LEGENDS

**eTable1 | Chr1q gain associated DEGs robust to cell type or in excitatory neurons, inhibitory neurons, and astrocytes.** Chr1q gain associated DEGs robust to cell type were identified using limma (see eMethods) whereas chr1q gain associated DEGs in excitatory neurons, inhibitory neurons, and astrocytes were identified using Seurat::FindMarkers (see eMethods). For genes located on chr1q, P values were corrected for multiple testing with Holm’s method; for genes not located on chr1q, the Benjamini-Hochberg method was used. Column names in the first tab: ‘gene’ - HGNC gene symbol of the DEG, ‘logFC’ - average log2-fold change, ‘AveExpr’ - average log-scaled expression score of the DEG (see eMethods), ‘t’ - value of t statistic from limma, ‘P.Value’ - unadjusted P value from limma, ‘adj.P.Val’ - Benjamini-Hochberg adjusted P value. Column names in remaining tabs: ‘gene’ - HGNC gene symbol of the DEG, ‘p_val’ - unadjusted P value, ‘avg_diff’ - average log2-fold change, ‘pct.1’ - percent of cells with chr1q gain in which the DEG was detected, ‘pct.2’ - percent of cells without chr1q gain in which the DEG was detected, ‘p_val_adj’ - Benjamini-Hochberg adjusted P value.

**eTable2 | Enrichment analysis of chr1q gain associated DEGs robust to cell type or in excitatory neurons, inhibitory neurons, and astrocytes.** Enrichment analysis was performed separately on upregulated DEGs and downregulated DEGs. Column names: ‘Library’ - gene set library, ‘Term’ - name of the gene set within the gene set library, ‘Overlap’ - number of DEGs overlapping with the gene set over the number of genes within the gene set, ‘P.value’ - unadjusted *P* value from Fisher’s exact test, ‘Odds.Ratio’ - odds ratio from Fisher’s exact test, ‘Genes’ - the DEGs overlapping with the gene set, ‘Combined.Score’ - a score that combines the odds ratio and *P* value, ‘DE_direction’ - the direction of differential expression of the DEGs.

**eTable 3 | Excitatory neuron, inhibitory neuron, and astrocyte subcluster DEGs.** Column names: ‘gene’ - HGNC gene symbol of the DEG, ‘p_val’ - unadjusted *P* value, ‘avg_diff’ - average log_2_-fold change, ‘pct.1’ - percent of cells in the subcluster of interest in which the DEG was detected, ‘pct.2’ - percent of cells not in the subcluster of interest in which the DEG was detected, ‘p_val_adj’ - Benjamini-Hochberg adjusted *P* value, ‘cluster’ - the subcluster of interest.

**eTable 4 | Enrichment analysis of excitatory neuron, inhibitory neuron, and astrocyte subcluster DEGs.** Enrichment analysis was performed separately on upregulated DEGs and downregulated DEGs. Column names: ‘Library’ - gene set library, ‘Term’ - name of the gene set within the gene set library, ‘Overlap’ - number of DEGs overlapping with the gene set over the number of genes within the gene set, ‘P.value’ - unadjusted *P* value from Fisher’s exact test, ‘Odds.Ratio’ - odds ratio from Fisher’s exact test, ‘Genes’ - the DEGs overlapping with the gene set, ‘Combined.Score’ - a score that combines the odds ratio and *P* value, ‘DE_direction’ - the direction of differential expression of the DEGs.

**eFigure 1.**
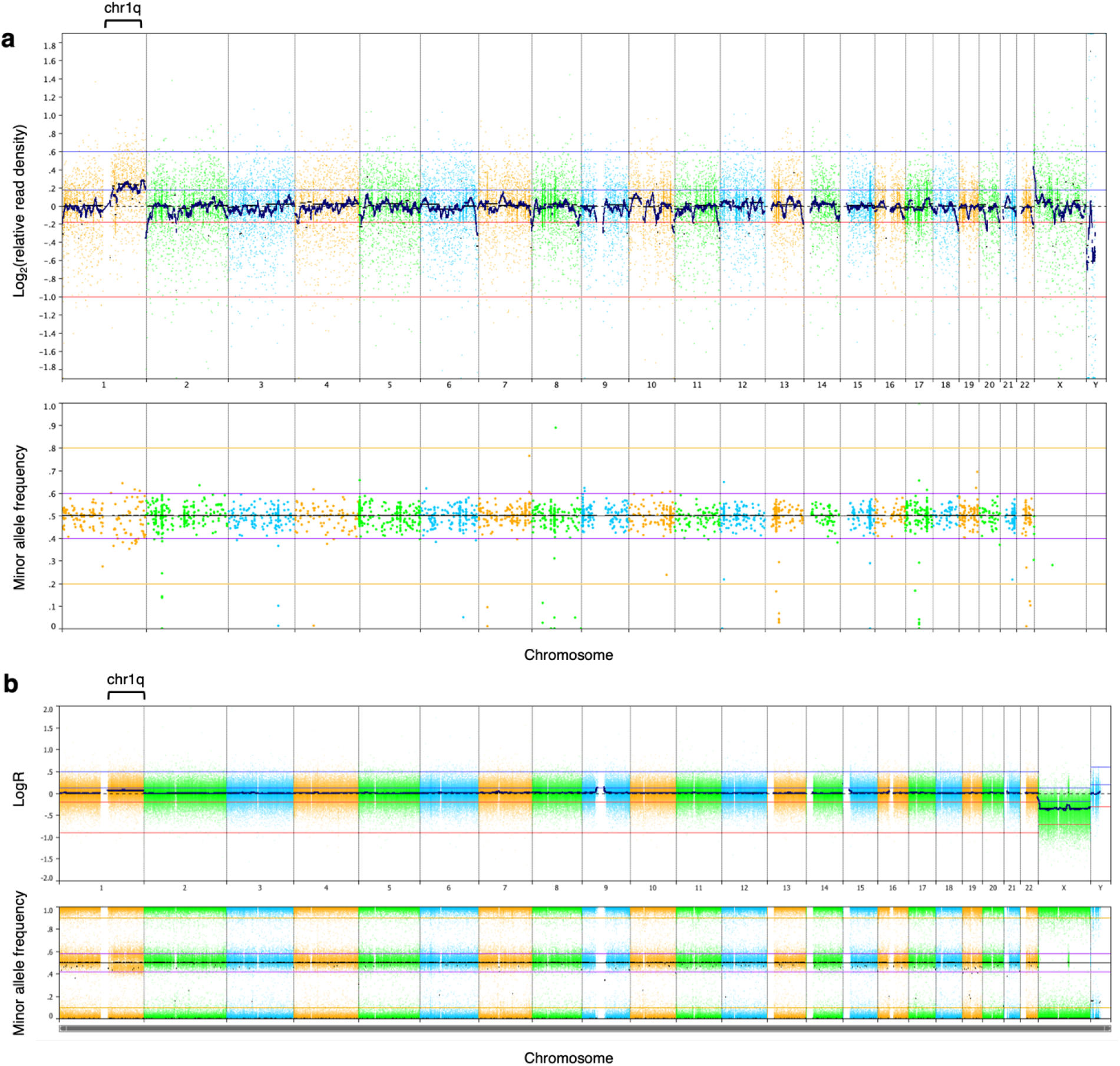
UCSF500 read density across all chromosomes and additional characterization of mosaic chr1q gain with high density SNP array. **a**, Log_2_-scaled relative read density (top) and minor allele frequency (bottom) from the UCSF 500 Cancer Gene Panel data shown in Fig. 1, shown for all chromosomes. **b**, LogR values (top, see eMethods) and minor allele frequency (bottom) from the Illumina CytoSNP 850 K array applied to genomic DNA extracted from neuronal nuclei isolated by fluorescence-activated nuclei sorting from fresh frozen resected brain tissue from the patient’s second surgery at age 3.

**eFigure 2.**
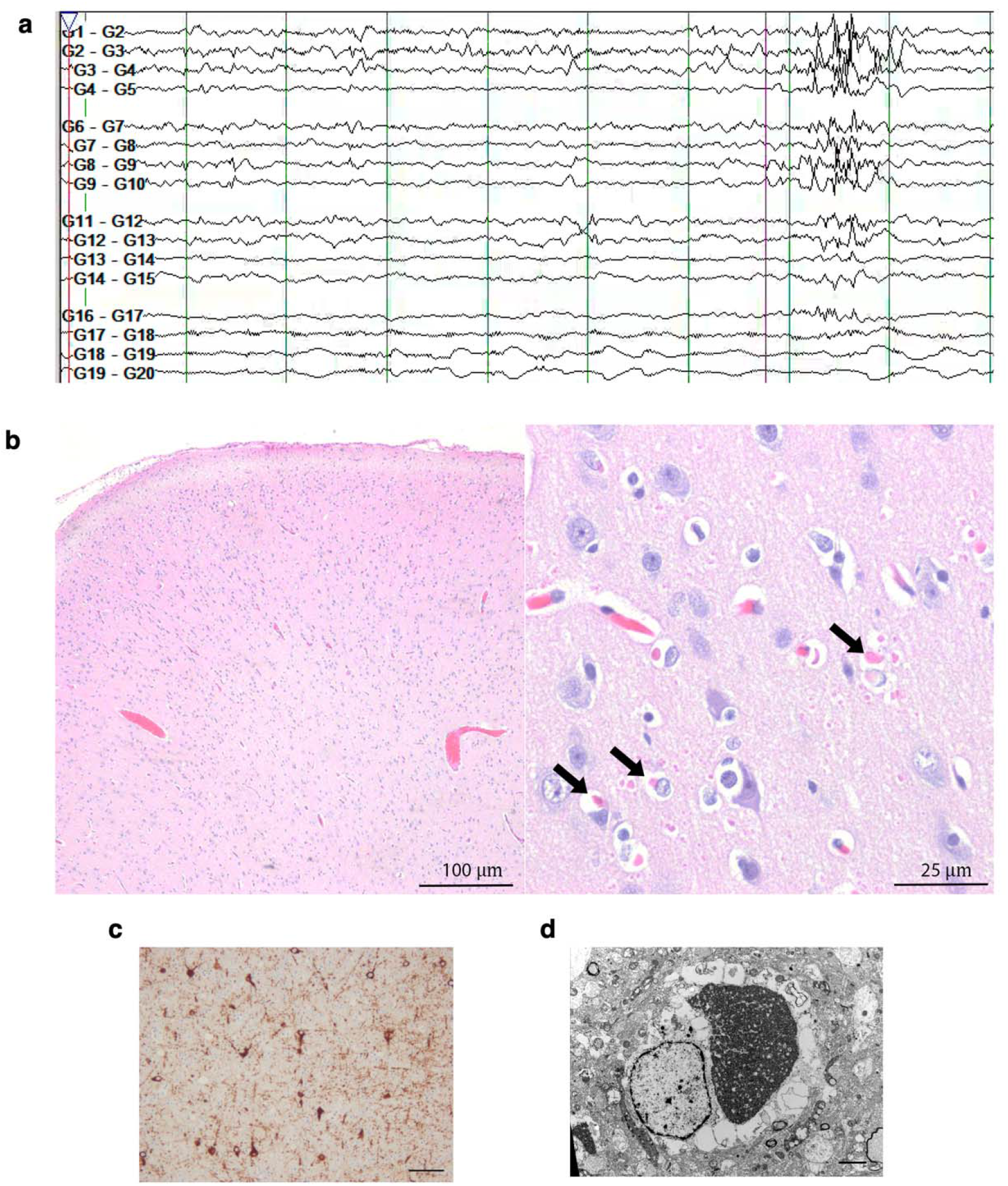
Additional clinical and neuropathological characterization. **a**, Intra-operative electrocorticography recording from the surgery at age 3 years demonstrating right frontal localization of epileptiform activity; the electrode array (4x5 grid, 20 electrodes) was placed over the right frontal lobe (see eMethods in Supplemental Information). **b**, Low (left) and high (right) power magnification hematoxylin and eosin (H&E) images from the cortex, showing relatively well-preserved laminar pattern. High power magnification reveals numerous bright eosinophilic inclusions within the cytoplasm of glial and neuronal cells (black arrows). Smaller inclusions can also be seen in the neuropil. The inclusions were in various sizes and shapes but typically attained spherical forms, particularly within the cytoplasm. **c**, MAP2 immunohistochemistry of white matter in resected cortical tissue at age 3 years demonstrating numerous heterotopic neurons. Scale bar, 100 µm. **d**, Electron microscopy of resected cortical tissue at age 5 years demonstrating an electron-dense cytoplasmic inclusion in an astrocyte. Scale bar, 2 µm.

**eFigure 3.**
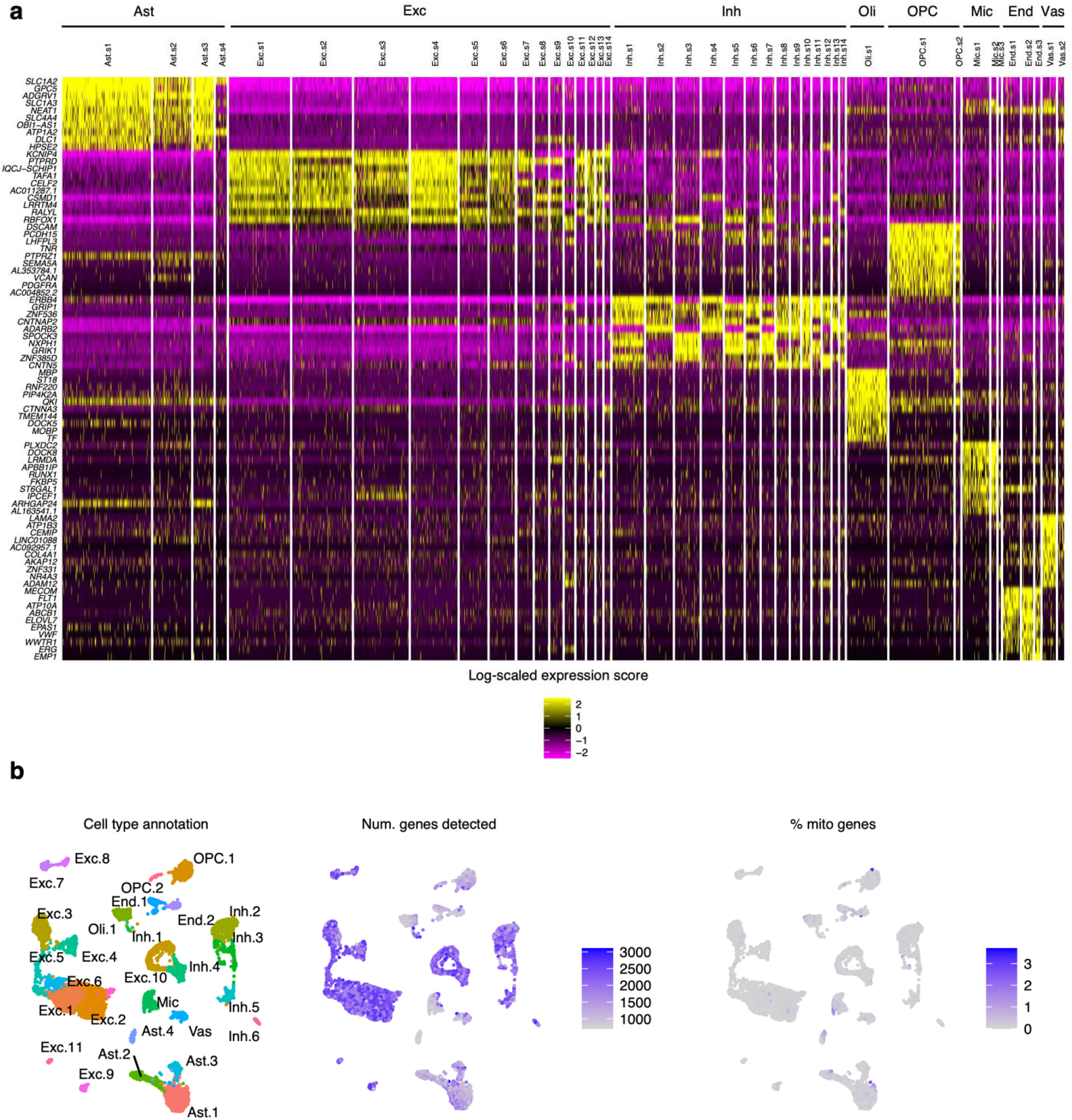
Appropriate expression of cell-type specific markers and expected variation in median number of genes detected seen on quality-control analysis. **a**, Heatmap of the top 10 cell type-specific genes across all major cell types captured by snRNA-seq, with cells organized by subcluster assignment. **b**, UMAP of all cells recovered from snRNA-seq colored by cluster assignment, median number of genes detected, or percent of genes from the mitochondrial genome.

**eFigure 4.**
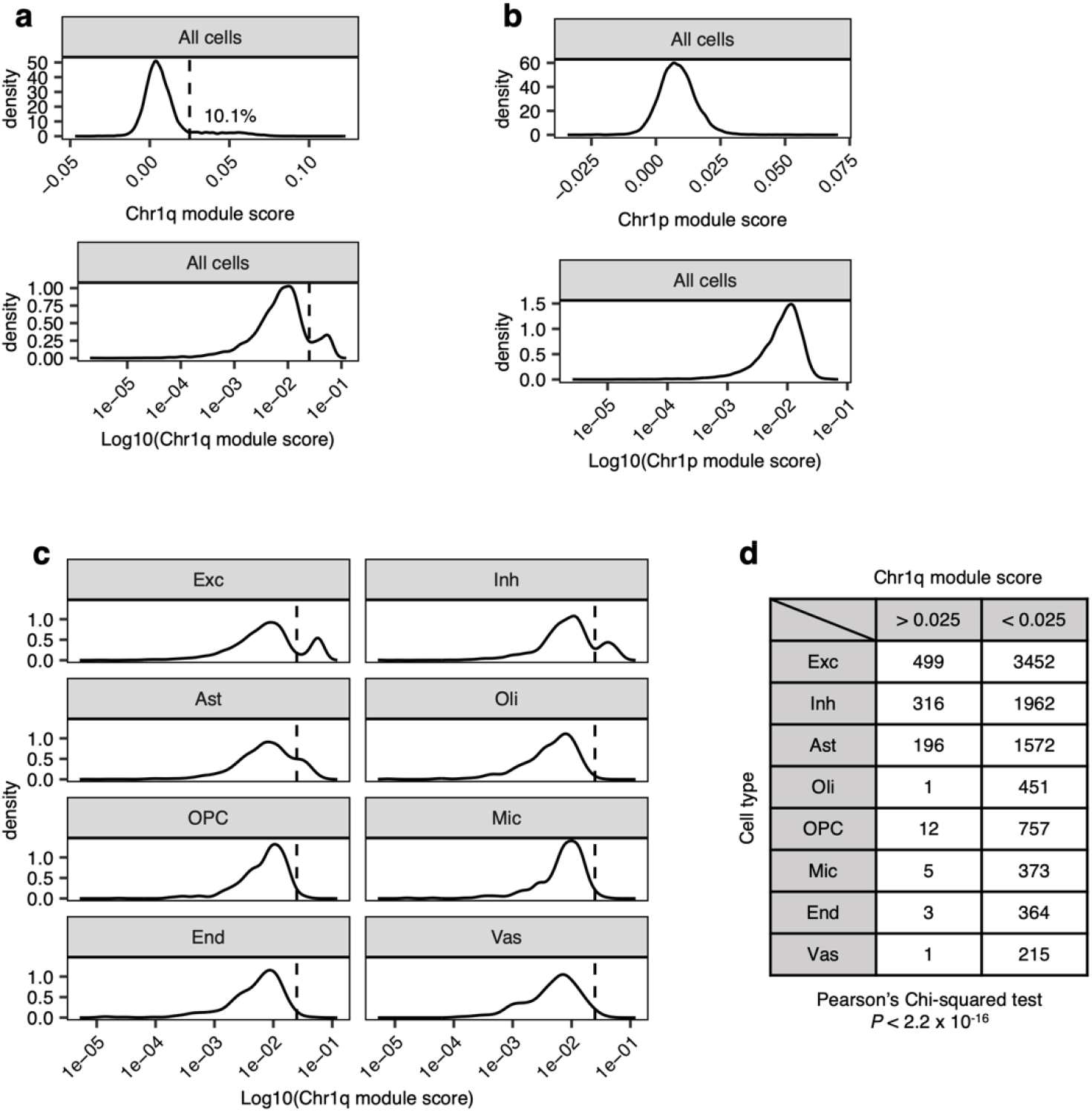
Inference of chr1q gain via chr1q module score. **a**, Density plot of the distribution of chr1q module scores across all cells, plotted on a linear (top) or logarithm (bottom) scale, with the cutoff value of 0.025 highlighted as a dashed vertical line. **b**, Density plot of the distribution of chr1p module score across all cells, plotted on a linear (top) or logarithmic (bottom) scale. **c**, Density plots of the distribution of chr1q module scores in each cell type, in logarithmic scale. **d**, Contingency table of number of cells with (chr1q module score > 0.025) vs without (chr1q module score < 0.025) inferred chr1q gain in each cell type, with the *P* value from Pearson’s Chi-squared test shown below.

**eFigure 5.**
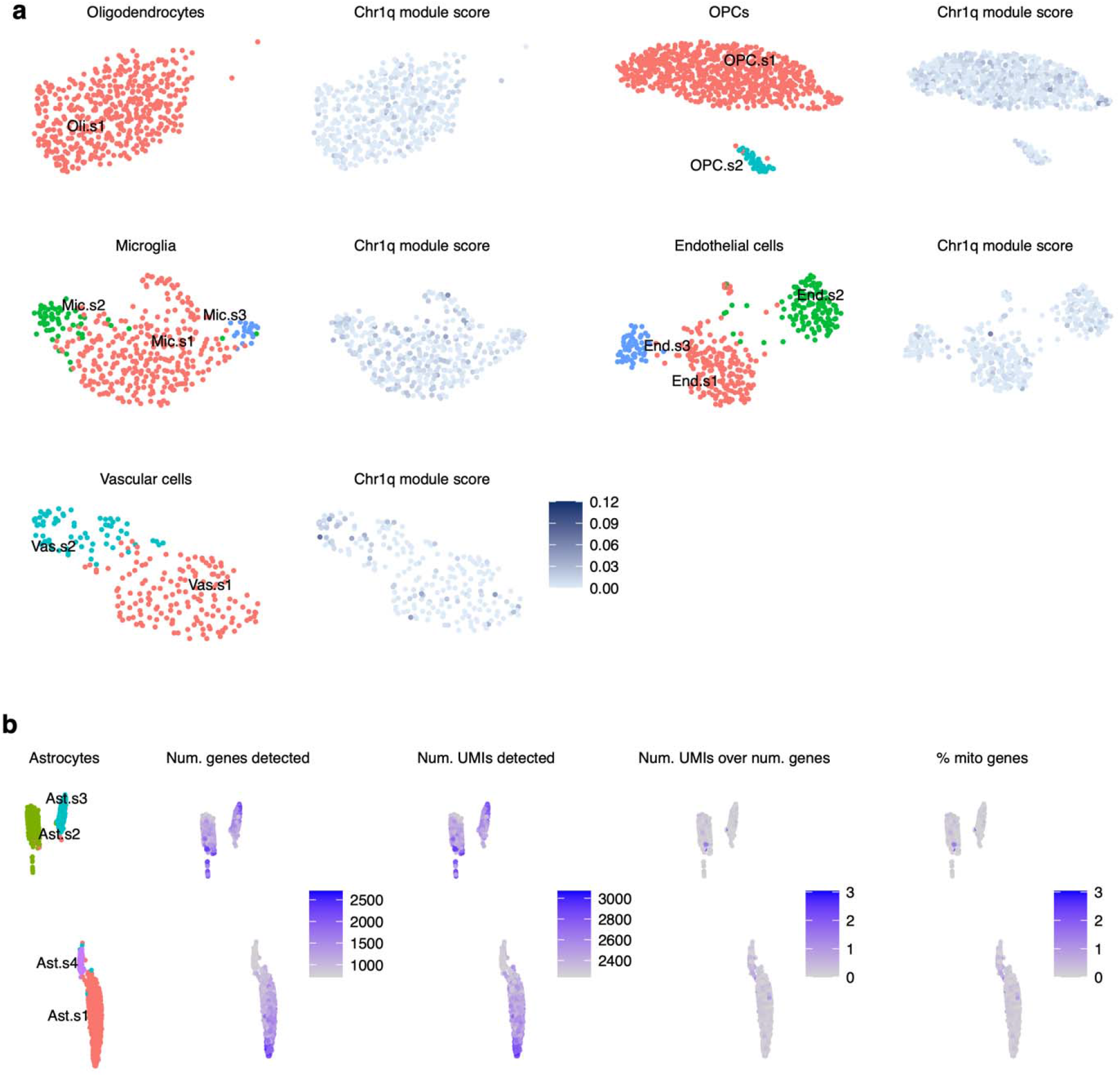
Subpopulations of OPCs, microglia, endothelial cells, and vascular cells seen on subclustering of cell types without inferred chr1q gain; low median number of genes detected in Ast.s4 seen on subclustering of astrocytes. **a**, UMAP of cell types without inferred chr1q gain colored by subcluster assignment or chr1q module score. **b**, UMAP of astrocytes colored by subcluster assignment, median number of genes or UMIs detected, the ratio of the number of detected UMIs to genes, or percent of genes from the mitochondrial genome. The low median number of genes detected in Ast.s4 makes it unlikely that Ast.s4 corresponds to astrocyte-neuron doublets, which tended to have >2000 median number of genes detected and were removed during quality control.

**eFigure 6.**
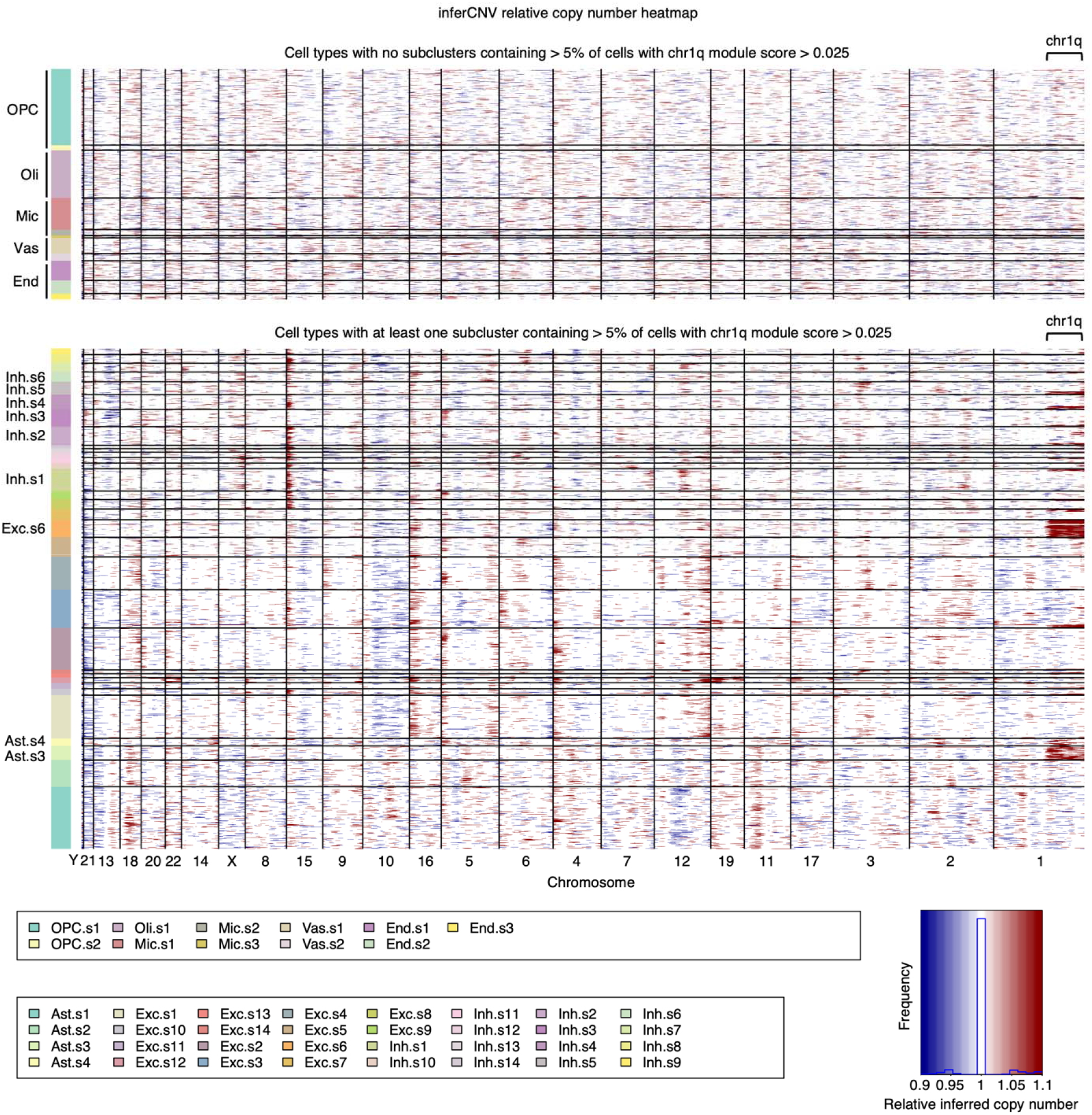
Inference of chromosomal copy number using inferCNV. Heatmap of inferred relative chromosomal copy number values in cell types with no subclusters containing cells with inferred chr1q gain (top) vs cell types with at least one subcluster containing cells with inferred chr1q gain (bottom) based on the chr1q module score thresholding shown in eFigure 4.

**eFigure 7.**
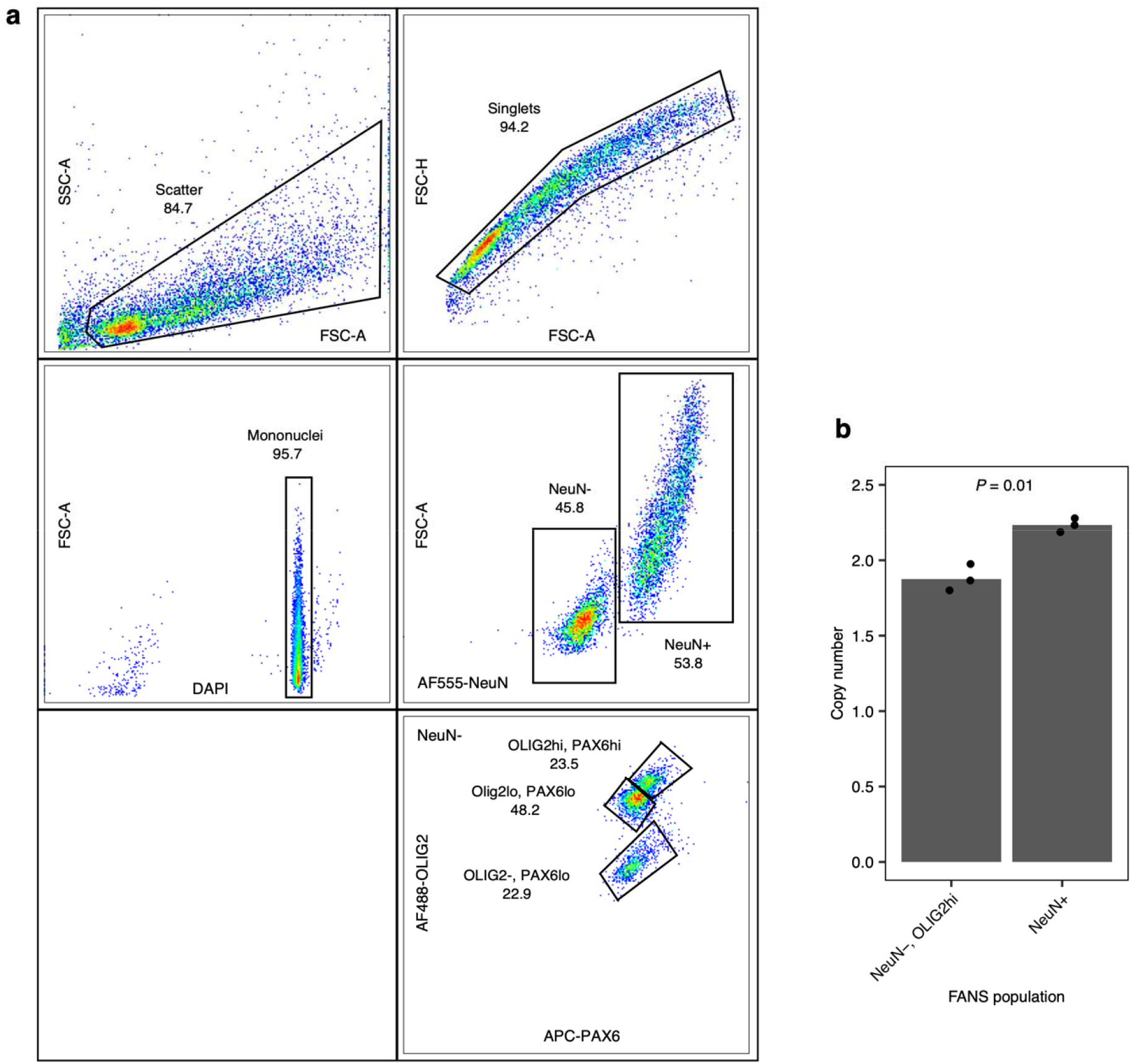
Validation of the cell-type specificity of mosaic chr1q gain using fluorescence activated nuclei sorting (FANS) and RT-PCR. **a**, FANS gating strategy. **b**, Chr1q copy number determined by RT-PCR in putative neurons (NeuN+) vs OPCs (NeuN-, OLIG2hi); technical replicates shown; *P* value calculated using two-sided Student’s t test.

**eFigure 8.**
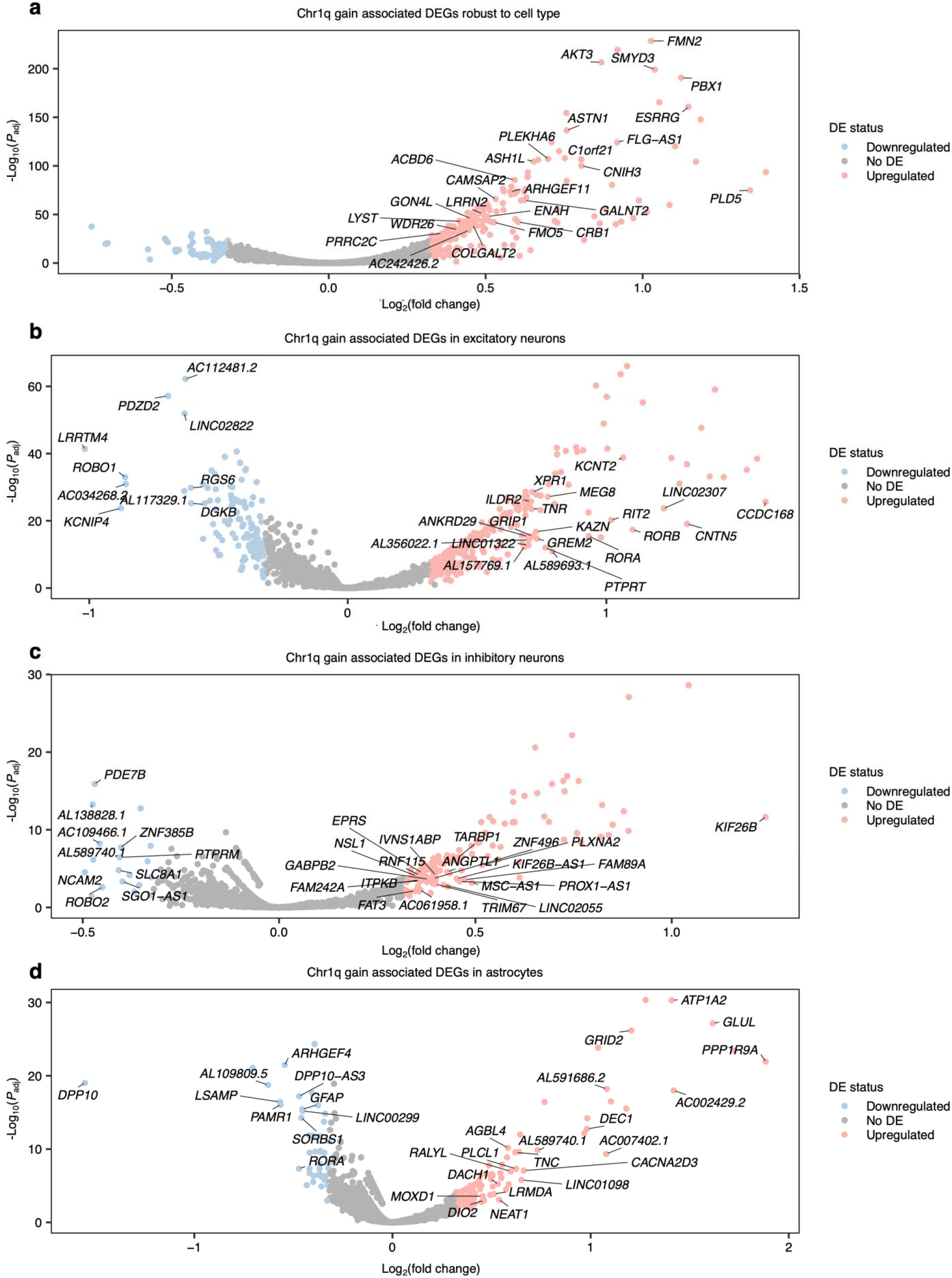
Cell type-robust and cell type-specific chr1q gain-associated DEGs. Volcano plots of chr1q gain associated DEGs robust to cell type (**a**), in excitatory neurons (**b**), inhibitory neurons (**c**), or astrocytes (**d**). DEGs shared among excitatory neurons, inhibitory neurons, and astrocytes are marked in panel **a**; DEGs specific to excitatory neurons, inhibitory neurons, or astrocytes are marked in panels **b**, **c**, and **d**, respectively.

**eFigure 9.**
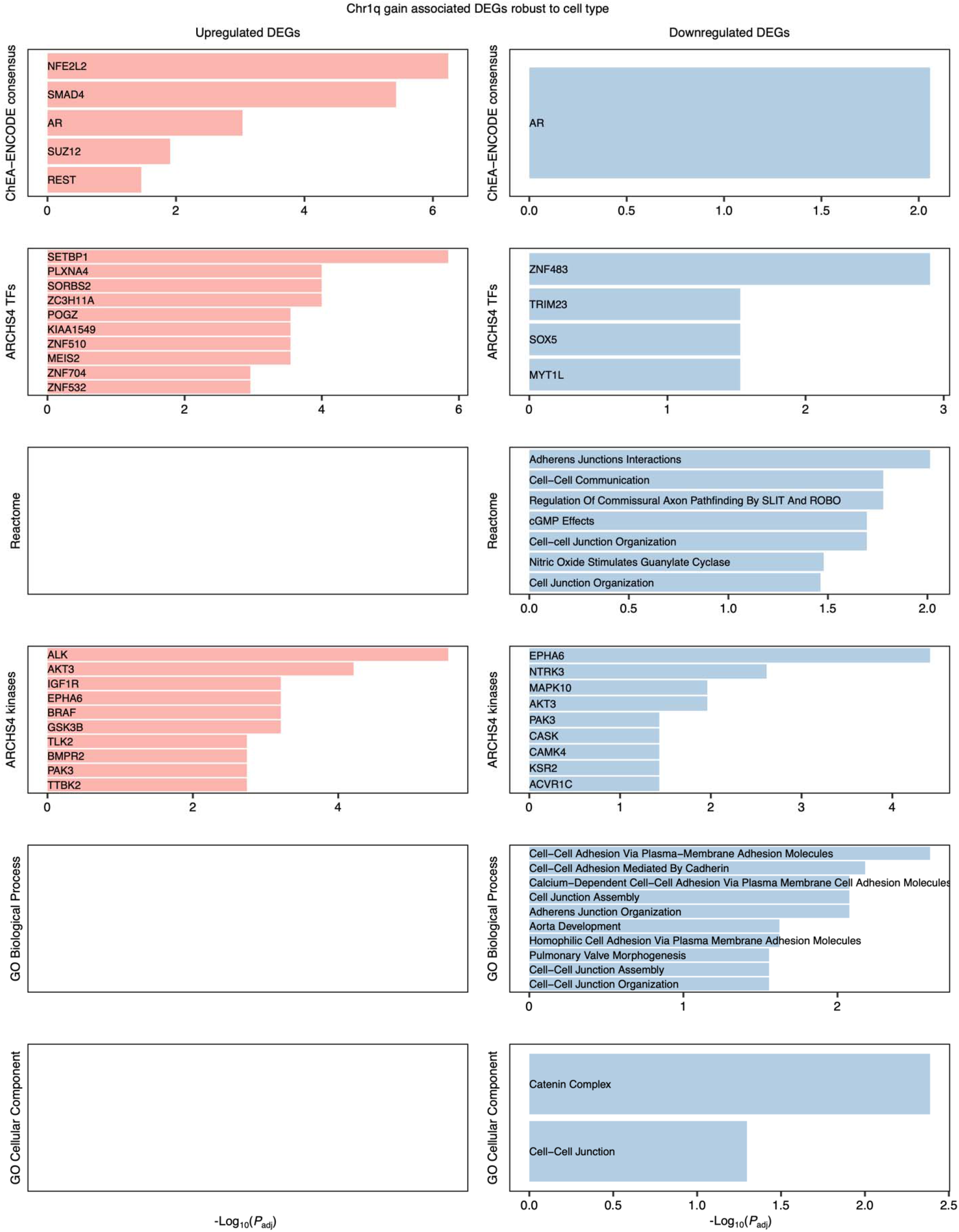
Enrichment analysis of DEGs associated with chr1q gain robust to cell type. See eMethods for details regarding the gene set libraries included for analysis. Only terms with adjusted *P* value < 0.1 are shown.

**eFigure 10.**
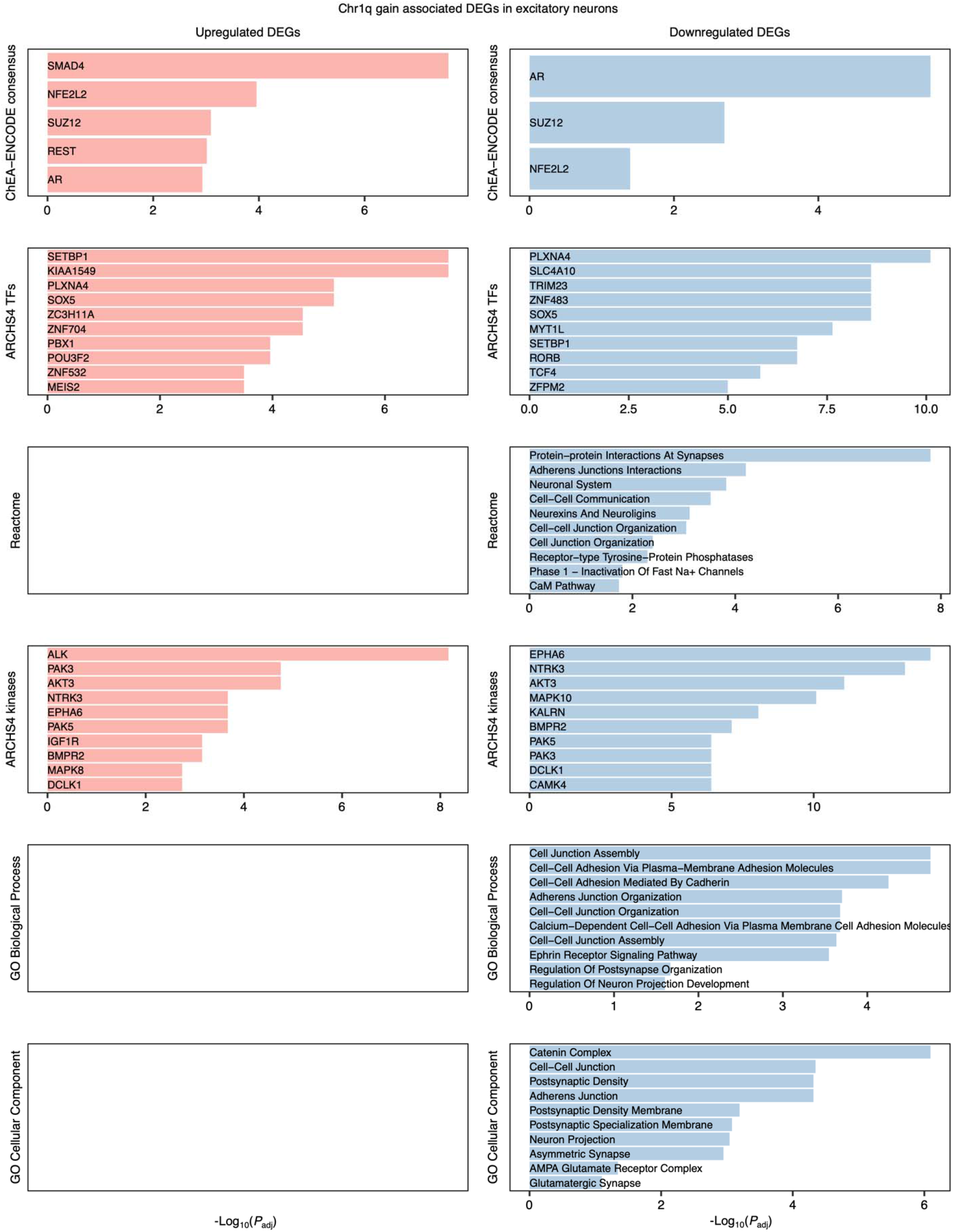
Enrichment analysis of DEGs associated with chr1q gain in excitatory neurons. See eMethods for details regarding the gene set libraries included for analysis. Only terms with adjusted *P* value < 0.1 are shown.

**eFigure 11.**
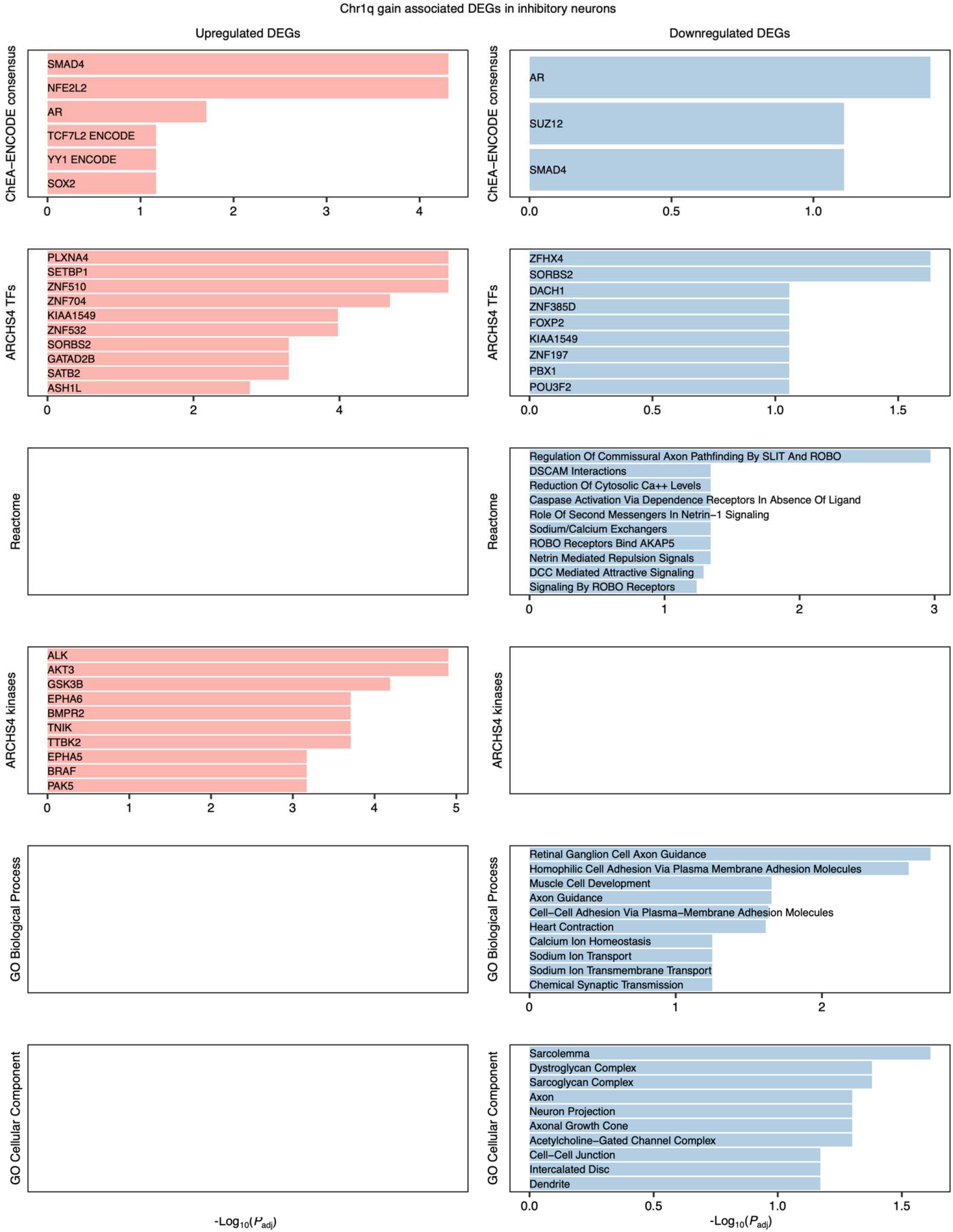
Enrichment analysis of DEGs associated with chr1q gain in inhibitory neurons. See eMethods for details regarding the gene set libraries included for analysis. Only terms with adjusted *P* value < 0.1 are shown.

**eFigure 12.**
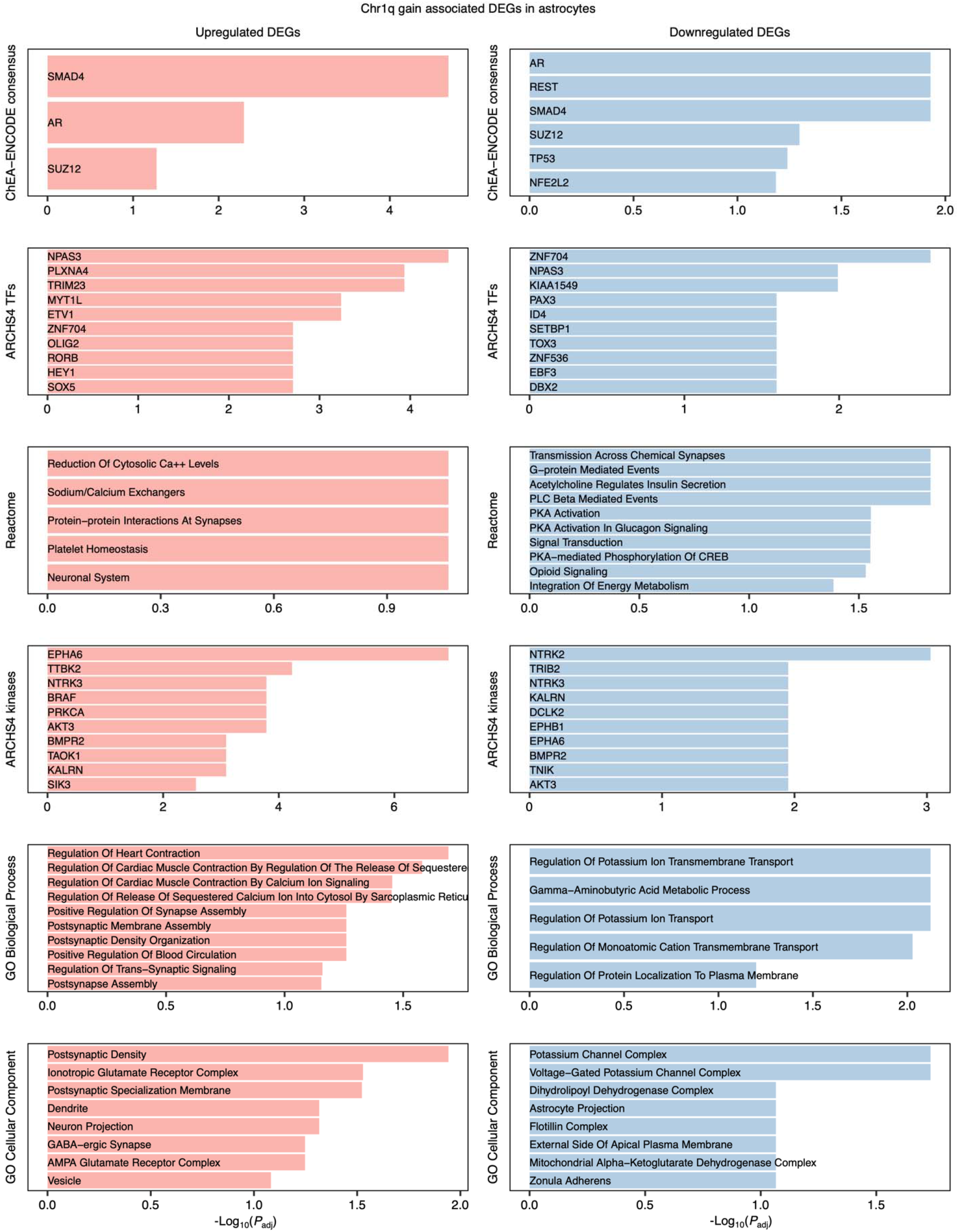
Enrichment analysis of DEGs associated with chr1q gain in astrocytes. See eMethods for details regarding the gene set libraries included for analysis. Only terms with adjusted *P* value < 0.1 are shown.

**eFigure 13.**
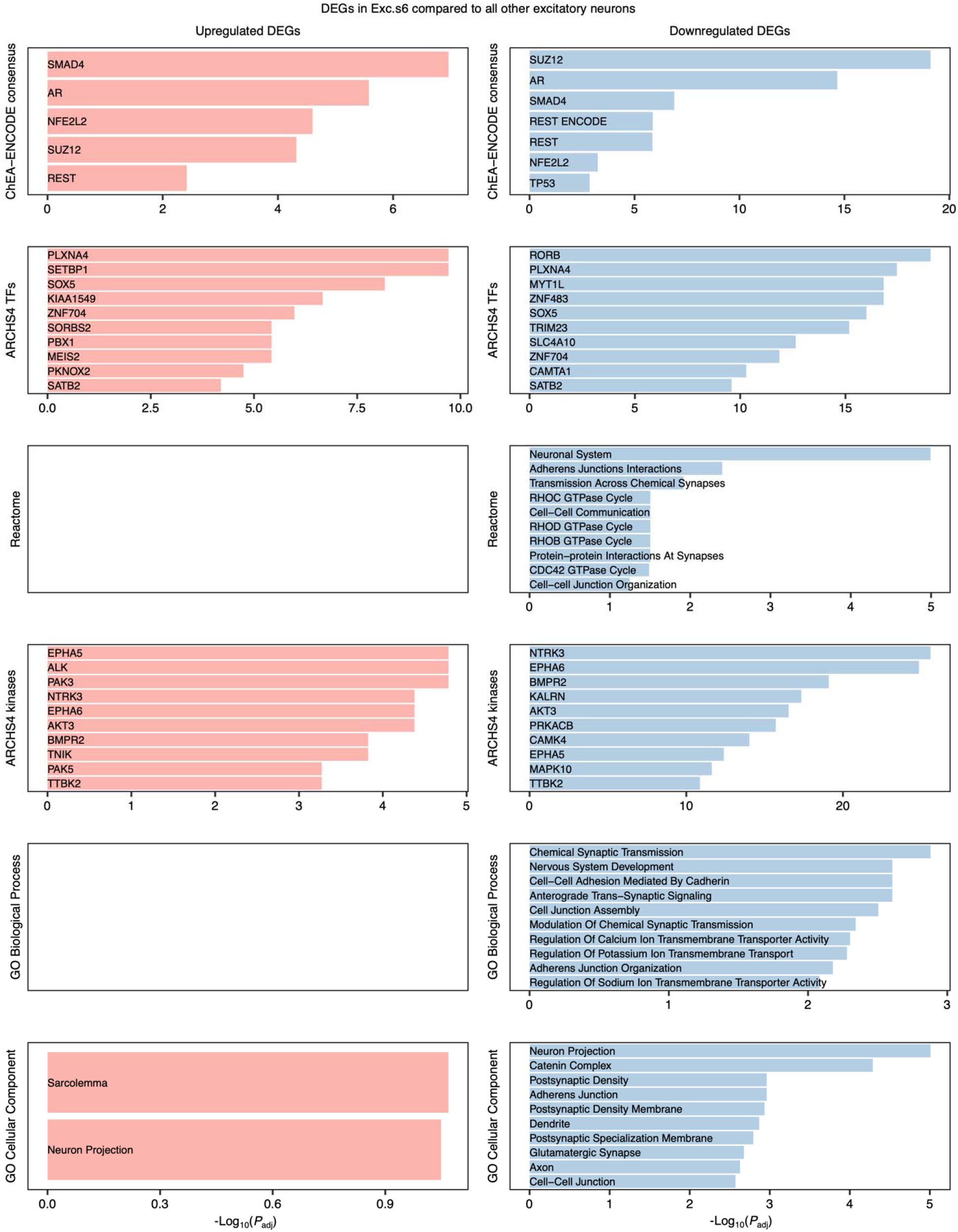
Enrichment analysis of DEGs in Exc.s6 compared to all other excitatory neurons. See eMethods for details regarding the gene set libraries included for analysis. Only terms with adjusted *P* value < 0.1 are shown.

**eFigure 14.**
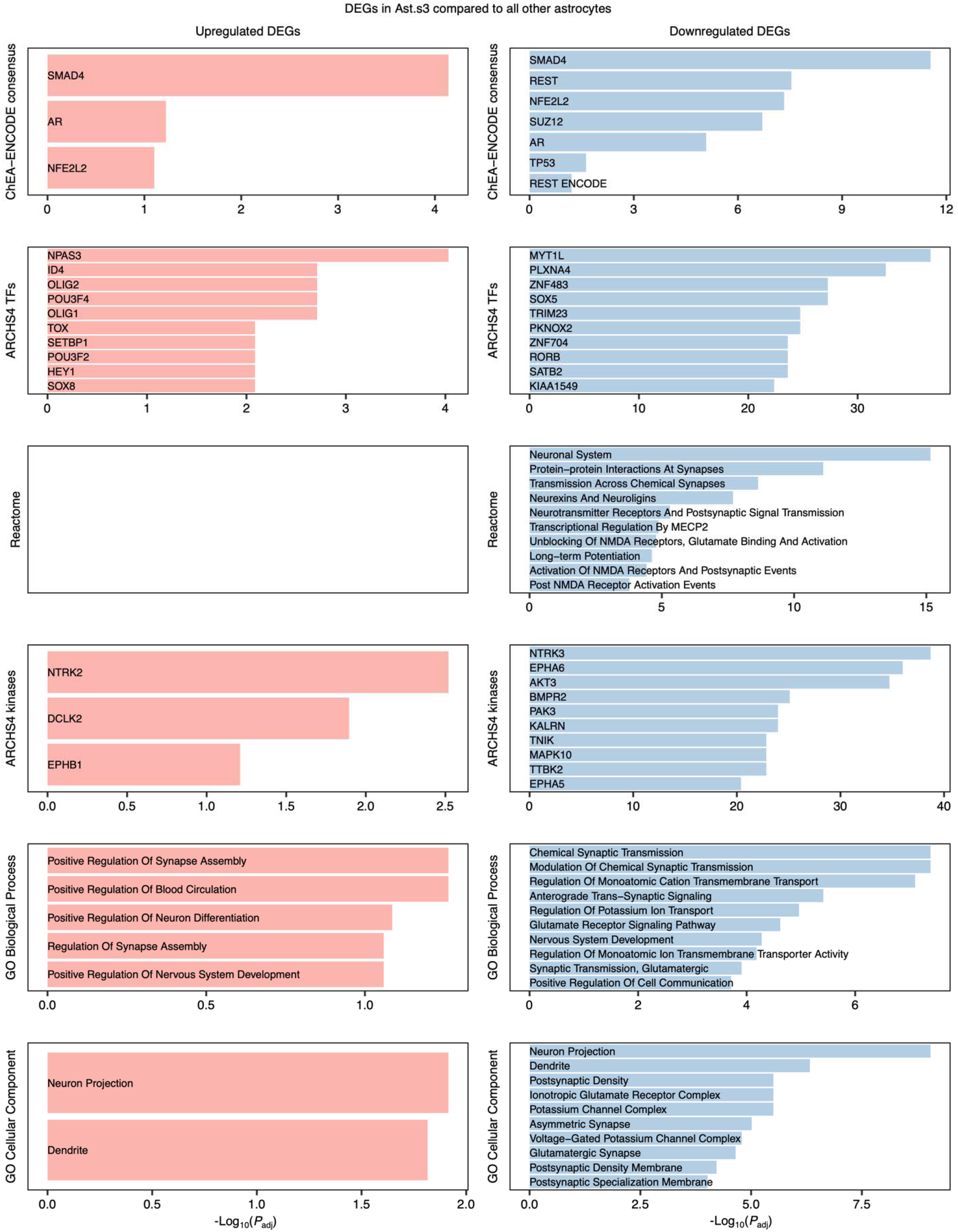
Enrichment analysis of DEGs in Ast.s3 compared to all other astrocytes. See eMethods for details regarding the gene set libraries included for analysis. Only terms with adjusted *P* value < 0.1 are shown.

**eFigure 15.**
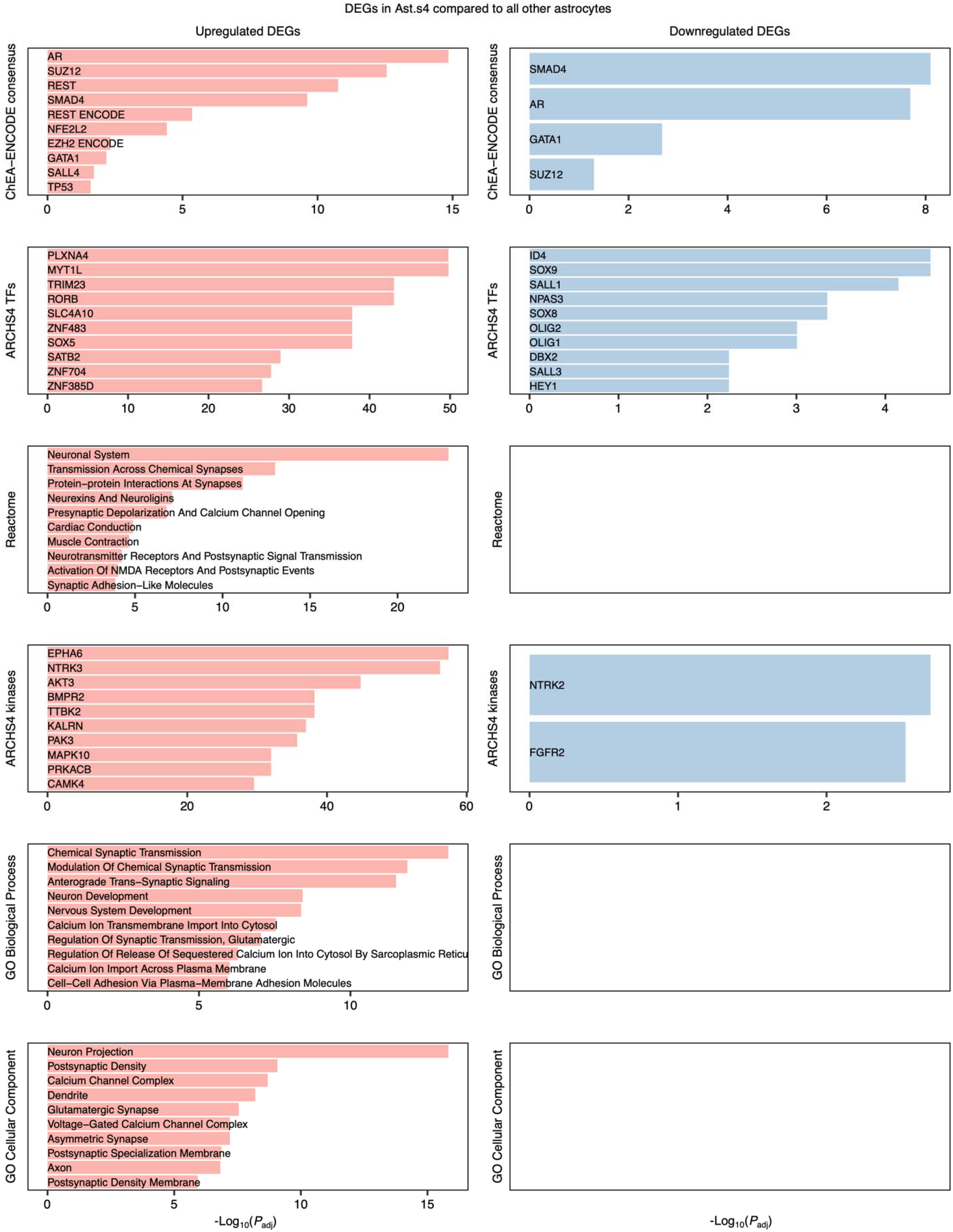
Enrichment analysis of DEGs in Ast.s4 compared to all other astrocytes. See eMethods for details regarding the gene set libraries included for analysis. Only terms with adjusted *P* value < 0.1 are shown.

**eFigure 16.**
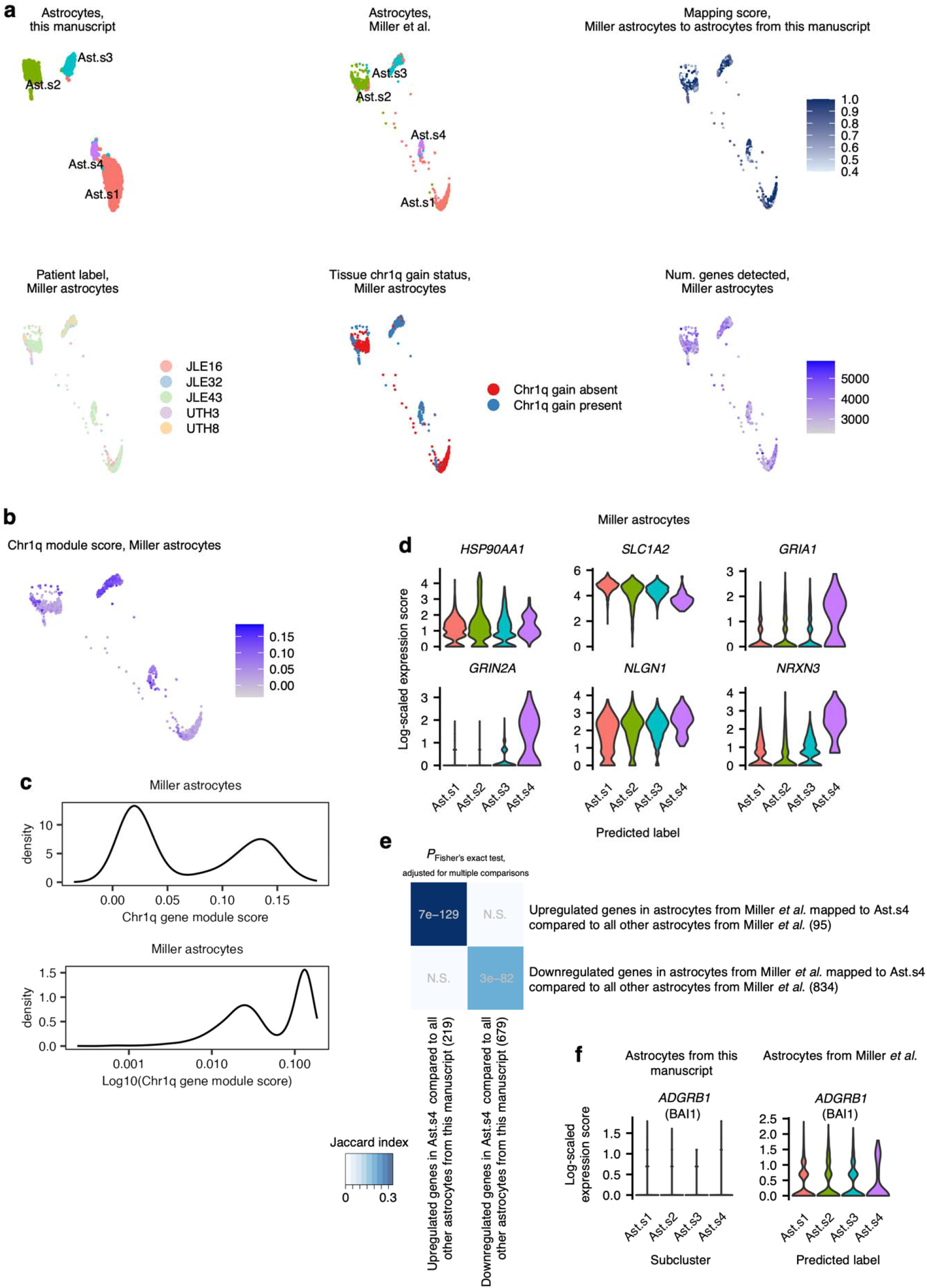
Label-transfer analysis of snRNA-seq data from additional HPA cases with mosaic chr1q gain from Miller *et al.* **a**, UMAP of astrocytes from this manuscript (top left) used as a reference for projecting astrocytes from Miller *et al.* (top middle), with the quality of the cross-dataset mapping shown as a mapping score (top right). The patient label (bottom left) and chr1q gain status of the tissue from which the cells were derived (bottom middle) as well as QC metrics such as the number of genes detected (bottom right) of astrocytes from Miller *et al.* are also shown. **b**, chr1q module score in astrocytes from Miller *et al.* visualized on the same UMAP embedding as in **a**. **c**, Density plot of the distribution of chr1q module scores across astrocytes from Miller *et al.*, plotted on a linear (top) or logarithm (bottom) scale. **d**, Log-scaled expression of selected synapse-associated transcripts in astrocytes from Miller *et al.*, grouped by the astrocyte subcluster in this manuscript to which they are mapped (predicted label). **e**, Heatmap of the Jaccard index (size of the intersection divided by the size of the union of two sets) corresponding to the overlap between upregulated or downregulated genes in Ast.s4 compared to all other astrocytes from this manuscript or Miller *et al.* astrocytes mapped to Ast.s4 compared to all other Miller *et al.* astrocytes, overlaid by the Fisher’s exact test *P* value (adjusted for multiple comparisons). **f**, Log-scaled expression of *ADGRB1*, which encodes BAI1, in astrocytes from this manuscript (left) or astrocytes from Miller *et al*. (right).

